# On-rate modulation of cadherin interactions by chemical fragments

**DOI:** 10.1101/2020.08.30.274647

**Authors:** Akinobu Senoo, Sho Ito, Satoru Nagatoishi, Yutaro Saito, Go Ueno, Kouhei Yoshida, Takumi Tashima, Shota Kudo, Shinsuke Sando, Kouhei Tsumoto

**Affiliations:** Department of Chemistry and Biotechnology, Graduate School of Engineering, The University of Tokyo, 7-3-1, Hongo, Bunkyo-ku, Tokyo 113-8656, Japan; Graduate School of Life Science, University of Hyogo, 3-2-1 Kouto, Kamigori-cho, Ako-gun, Hyogo, 678-1297, Japan; ROD (Single Crystal Analysis) Group, Application Laboratories, Rigaku Corporation, 3-9-12 Matubara-cho, Akishima, Tokyo 196-8666, Japan; Institute of Medical Science, The University of Tokyo, 4-6-1, Shirokanedai, Minato-ku, Tokyo 108-8639, Japan; RIKEN SPring-8 Center, 1-1-1, Kouto, Sayo-cho, Sayo-gun, Hyogo 679-5148, Japan; Department of Bioengineering, Graduate School of Engineering, The University of Tokyo, 7-3-1, Hongo, Bunkyo-ku, Tokyo 113-8656, Japan

## Abstract

Many cadherin family proteins are associated with diseases such as cancer. Since cell adhesion requires homodimerization of cadherin molecules, a small-molecule regulator of dimerization would have therapeutic potential. Herein, we describe identification of a P-cadherin-specific chemical fragment that inhibits P-cadherin-mediated cell adhesion. Although the identified molecule is a fragment compound, it binds to a cavity of P-cadherin that has not previously been targeted, indirectly prevents formation of hydrogen bonds necessary for formation of an intermediate called the X dimer and thus modulates the on-rate of X dimerization. Our findings will impact on a strategy for kinetic regulation of protein-protein interactions and stepwise assembly of protein complexes using small molecules.

## Introduction

Protein assembly underlies almost all of the biological processes^1^. There has been so much reported structural information on homomers and heteromers in various kinds of functional complexes^2^. Modulators of protein assembly formation have potential as drugs and as probes to investigate protein function. However, the fundamental interaction of the assembly formation; protein-protein interactions are difficult to regulate with small molecules for several reasons: large surface areas are usually involved in interactions between proteins, whereas the accessible surface areas of chemical ligands are small, there are generally no substantial grooves at the protein-protein interface, and there are few natural inhibitors of protein-protein interactions to guide ligand design^3–5^.

One of the protein assembly formation can be found in the process of forming cell adhesion by classical cadherin family proteins, calcium-dependent cell adhesive molecules. Depending on biological context, cadherins can play tumor-promoting roles in various tissues^6^. P-cadherin is a classical cadherin family protein. Its overexpression in some cancer tissues has been reported^7–11^, and promotes metastasis and proliferation^13–18^. As formation of cell aggregates blocks anoikis^8^, and P-cadherin-mediated signaling is important for cancer cell survival ^17^, inhibition of P-cadherin function is a potential anticancer strategy. Cell adhesion is achieved through *trans* homodimerization between cadherins on apposed cells and then *cis* clustering of cadherins on the same cell surface^18^ (Fig. 1a).

**Fig. 1.**
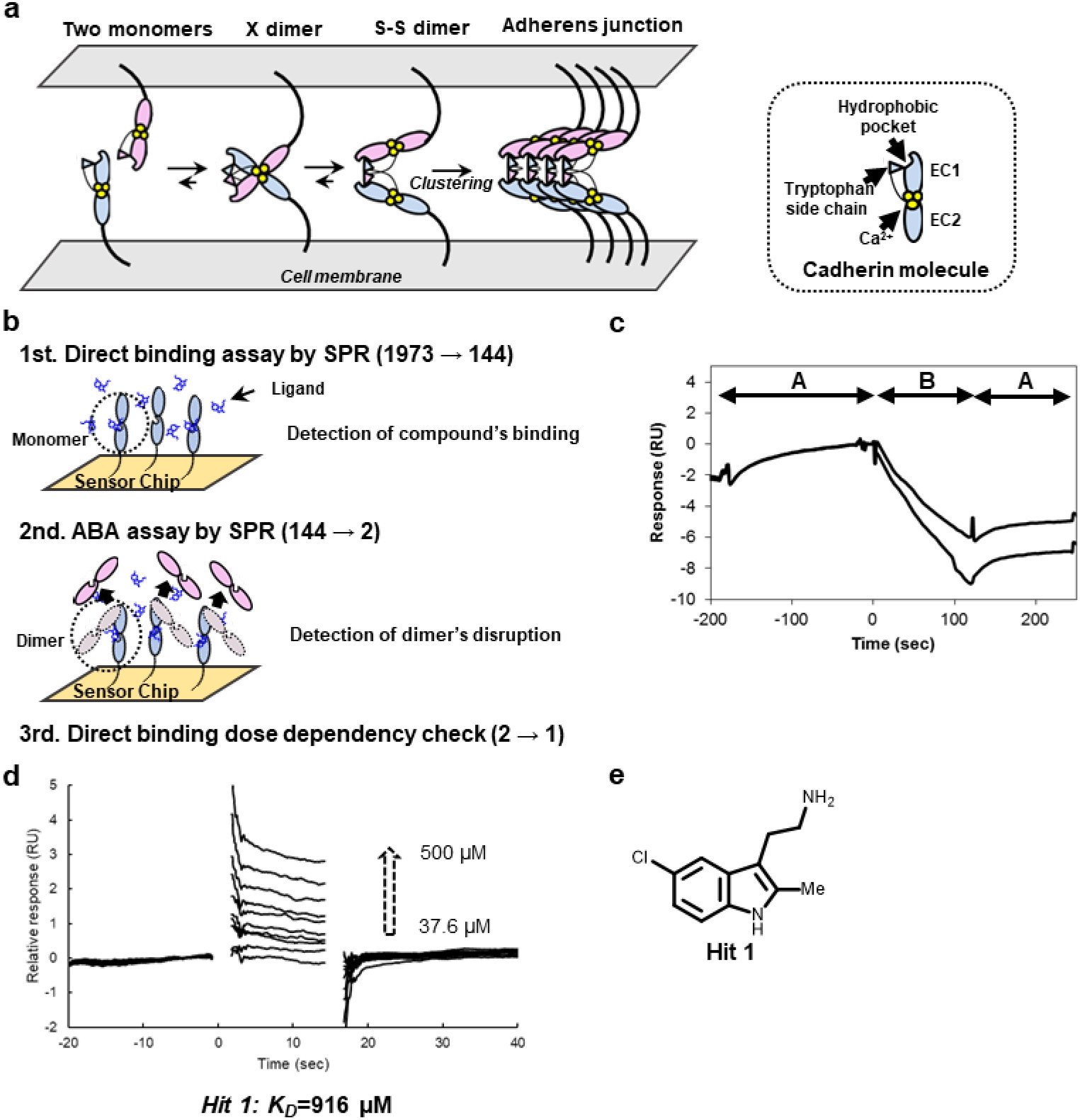
SPR-based fragment screening. (a) Illustration of the stepwise dimerization of P-cadherin to mediate cell adhesion and a schematic of the protein (EC12). (b) Schematic of fragment screening. (c) Representative sensorgram from ABA assay with EC12, in monomer-S-S dimer equilibrium, was immobilized on Sensor Chip CM5. The decrease of response upon injection of solution B shows that the compound disrupted the dimer formed on the sensor chip. (d) Dose-response analysis of binding of Hit 1 to REC12 immobilized on Sensor Chip SA. The putative *K_D_* was calculated by the Scatchard method using the SPR responses in equilibrium. Note that the *K_D_* value is not reliable, since the binding response was not saturated even at the highest concentration of Hit 1. (e) Chemical structure of Hit 1.

For some type I classical cadherins, including P-cadherin, *trans* homodimerization has been intensively studied, and the two extracellular domains of five on the protein that interact during the step-wise process have been identified^19–22^ (Fig. 1a). The X dimer is an intermediate state that favorably promotes the final dimerized state, which is called the strand-swap dimer (S-S dimer). The S-S dimer has a unique binding mode in which a tryptophan residue of the N terminal strand of one monomer is swapped into a hydrophobic pocket of the other monomer. Based on this binding mode, several peptide mimetic ligands have been reported^23–28^; however, no ligands that bind to the hydrophobic pocket have been identified, and the molecular basis of inhibition of homodimerization by these peptide mimetics has not been elucidated. Orthosteric inhibition of S-S dimerization, which is the last stage in the assembly process, may not be the best way to regulate the process. We have to change direction to find other way to effectively regulate the cell adhesive assembly with a small molecule.

In this study, we performed screen for inhibitors at any stage of the process. We screened a library that contained small-molecule drug-like fragments and identified a compound that bound to a novel ligand binding site unique to P-cadherin. The compound inhibited cell adhesion through the modulation of the on-rate of X dimerization, indirectly blocking formation of hydrogen bonds that stabilize the X dimer.

## Results

### SPR-based fragment screen

SPR-based fragment screen was performed as described previously^29^ (Fig. 1b). The fragment library was ordered from Drug Discovery Initiative; it contained 1973 compounds. As a primary screen, a direct binding assay was performed. Monomer mutant of P-cadherin called REC12^21^ was immobilized on the Sensor Chip SA via biotin-streptavidin capture method and 100 μM of each compound was injected onto the sensor chip surface. Based on immobilization level and molecular weights of REC12 and fragment compounds, we estimated that the R_MAX_ value of the binding response would be around 20 RU. Of the 1973 compounds in the library, 144 compounds had binding responses greater than 10 RU (Supplementary Fig. S1a).

As a secondary screen, we immobilized a construct called EC12 on the Sensor Chip CM5.This construct is in equilibrium between monomer and S-S dimer. We performed the so-called ABA assay, in which two solutions, A and B, are injected successively. In our experiments, 2 μM EC12 was used as a solution A, and 100 μM of each of the compounds identified in the primary screen was used as a solution B. In both A parts, the binding responses of monomer in the analyte and monomer covalently linked to the sensor chip surface were obtained. Two compounds significantly decreased the response in the B part (Supplementary Fig. S1b, Fig. 1c). These compounds bound to and disrupted the dimerization such that monomers that were not covalently immobilized on the sensor chip dissociated from the chip surface. Only one of the two compounds, Hit 1 here after, showed a dose-response dependency in the direct binding assay (Fig. 1d). The approximate *K_D_* value of Hit 1 (Fig. 1e) for the monomer mutant was calculated to be 916 μM.

### Identification of a ligand binding site for Hit 1

We employed X-ray crystallography to identify the binding site of Hit 1 to REC12 and to elucidate the inhibitory mechanism of Hit 1. REC12 was crystallized and soaked with Hit 1 at 10 mM final concentration. The resulting complex diffracted to 2.30Å resolution (Fig. 2a, Supplementary Table 1, Supplementary Fig. S2). Electron density corresponding to Hit 1 was observed inside a shallow cavity located between the EC1 and EC2 domains. Upon binding of Hit 1, the side chain of Y140 shifted relative to its position in the crystal structure of REC12 alone (PDB ID; 4zmz) (Fig. 2b), and Y140 and Hit 1 form a CH-π interaction. A water molecule was located in the cavity in the crystal structure of REC12 alone (PDB ID; 4zmz). Thus, the driving forces that result in of Hit 1 binding appear to be the interaction with Y140 residue, and the dehydration. Hit 1 also has some van der Waals interactions with R68, V98, T99, D100, and D137 (Fig.2c). In order to confirm the binding mode observed in the crystal structure, we performed a SPR-based direct binding assay using REC12 Y140R mutant. Hit 1 did not bind to this mutant (Supplementary Fig. S3), validating the contribution of Y140 to the binding observed in the crystal structure. Several residues around this binding pocket are reported to be important for X dimerization including Y140, D100 and Q101. In the X dimer, hydrogen bonds between Y140 and K14, and between Q101 and D100 are observed^21^. Although Hit 1 does not disrupt these hydrogen bonds directly, alternation of the structure of this region apparently inhibits X dimerization.

**Fig. 2.**
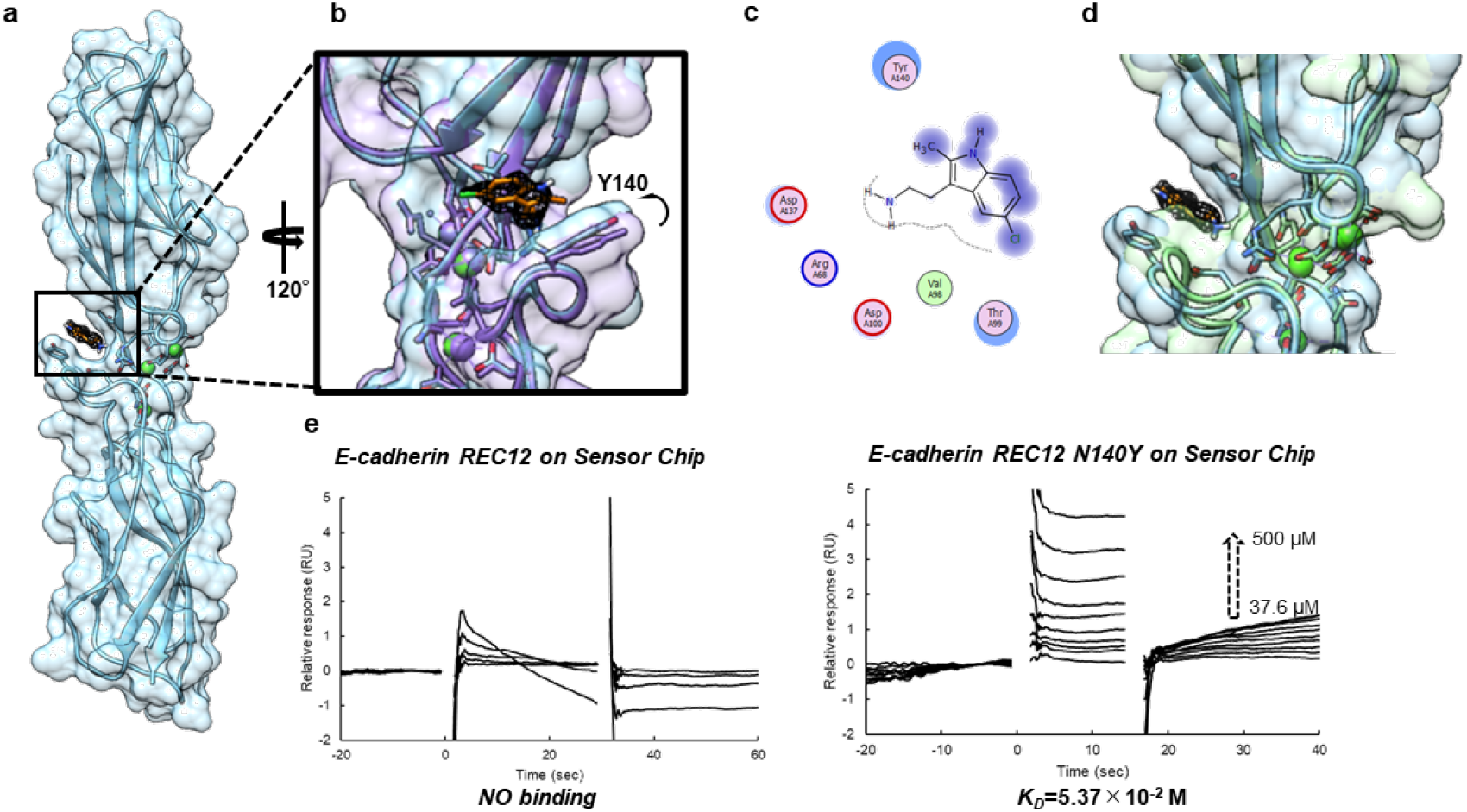
Hit 1 binds selectively to P-cadherin. (a) Structure of complex of REC12 with Hit 1. Hit 1 is colored orange. (b) Region of the Hit 1-binding cavity, illustrating the shift in the side chain of Y140 that occurs upon compound binding. 2mFo-DFc map at 1.0 σ level is shown. The P-cadherin monomer (PDB ID; 4zmz) is superposed (purple). (c) 2D interaction map of Hit 1 made with FLEV^32^. The reduction of solvent exposure upon Hit 1 binding is indicated by the halo-like disc around the residues. (d) Superposition of P-cadherin REC12-Hit 1 complex and E-cadherin structure (PDB ID 2o72; green). (e) Sensorgram from SPR-based binding assay of WT E-cadherin and E-cadherin N140Y with Hit 1. This experiment was repeated at least twice, and similar results were obtained.

Interestingly, when we superposed the P-cadherin-Hit 1 complex structure with that of E-cadherin (PDB; 2o72), we found that the volume of the cavity in E-cadherin was smaller than that in P-cadherin (Fig. 2d). Indeed, volumes of the cavities in the P-cadherin apo structure (PDB ID; 4zmz), the E-cadherin structure (PDB ID; 2o72), and the P-cadherin monomer-Hit 1 complex structure calculated using CASTp^30^ were 12.173 Å^3^, 5.441 Å^3^, 21.427 Å^3^, respectively, suggesting that P-cadherin but not E-cadherin, has a suitable cavity for the binding of Hit 1. Moreover, FTMap analysis^31^ regarded the region as a binding cavity only in the P-cadherin structure, not in E-cadherin structure (Supplementary Fig. S4). As expected, Hit 1 did not bind to E-cadherin but did bind to the E-cadherin N140Y mutant in the SPR-based direct binding assay (Fig. 2e). The N140Y point mutation did not affect the secondary structure of the recombinant E-cadherin, as confirmed using circular dichroism spectroscopy (Supplementary Fig. S5). Therefore, the binding cavity of Hit 1 could be used as a starting point for design of a ligand selective for P-cadherin.

### Effect of Hit 1 on X dimerization

To monitor the effect of Hit 1 on X dimerization, we used hydrogen-deuterium exchange mass spectrometry (HDX-MS). The method allowed us to monitor the extent to which the interface of X dimer is exposed to the solvent in the presence or in the absence of Hit 1. Pepsin treatment before LC-MS yielded peptide fragments from almost all regions of the P-cadherin molecule (Supplementary Fig. S6a, S6b. S6c). The HDX ratios for each peptide with and without Hit 1 were determined (Supplementary Fig. S6d and Fig. S6e). In the region that corresponds to one of the interfaces (residues 134-140 and 137-147) of MEC12, a construct that forms the X dimer but not the S-S dimer, the HDX ratio in the presence of Hit 1 was higher than in the absence of Hit 1, although not as high as that of REC12 (Fig. 3a). This result suggests that the equilibrium between monomer and X dimer shifted toward the monomer form in the presence of Hit 1. The assay does not reflect all the residues at the interface between monomers in the X dimer (Supplementary Fig. S6f, Fig. S6g). This may be because the interface of the N terminal region involved in the interaction of the X dimer is too dynamic to analyze in HDX-MS, since the HDX reaction occurs in regions exposed to the solvent due to dynamics.

**Fig. 3.**
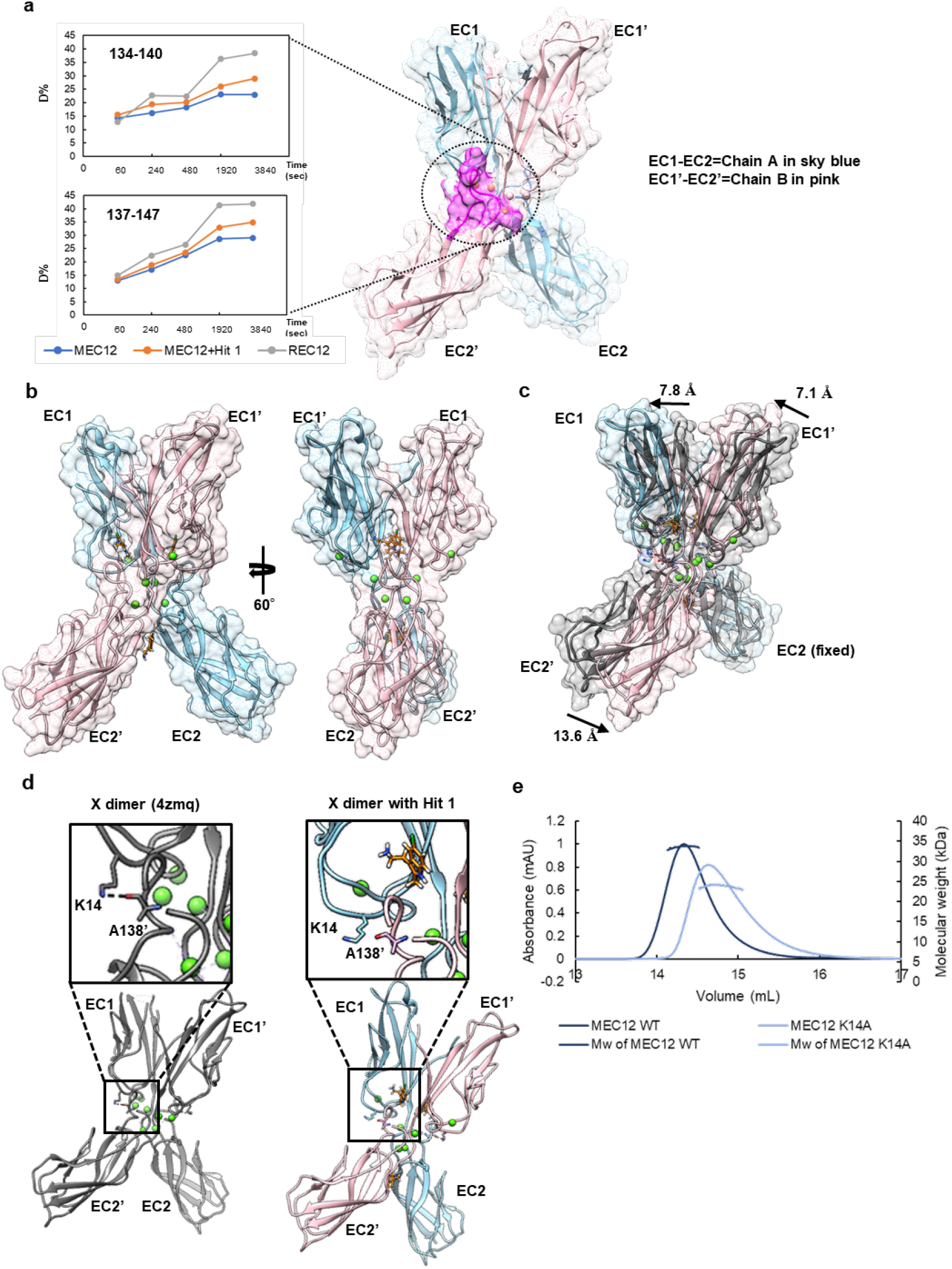
Effects on X dimerization by Hit 1. (a) Hydrogen-deuterium exchange ratio of two peptides 134-140 and 137-147 as a function of time. The region of these peptides is colored in magenta in the structure of MEC12, X dimer (PDB ID; 4zmq). Chain A of X dimer is colored in sky blue; chain B in pink. (b) Structure of the complex of the X dimer and Hit 1. Hit 1 (orange) is bound in the cavity around Y140 and at the intersection of EC2 domains. (c) Structural change caused by the binding of Hit 1. The angle of EC1 to EC2 is flatter compared to that in the apo X dimer (PDB ID; 4zmq, grey). The arrows indicate movement of domains. (d) Hydrogen bonds disrupted by the structural change that occurs upon Hit 1 binding. Left: Apo X dimer with K14-A138’ hydrogen bond is shown in dotted line. Right: Hit 1-bound X dimer; the K14-A138’ hydrogen bond is disrupted. Amino acid residues from chain A are shown in sky blue, those from chain B are in pink. (e) Size measurement using SEC-MALS. The WT X dimer trace is in dark blue, the K14A mutant trace is in light blue.

To further investigate how Hit 1 affects X dimerization, MEC12 was crystallized and soaked with Hit 1 at 10 mM final concentration, resulting in the complex structure at 2.45Å resolution (Fig. 3b, Supplementary Table 1). Three independent sets of electron density that can be modeled as Hit 1 were found, two of them in the cavity around Y140, the other at the intersection of EC2 domains (Fig. 3b). The Hit 1 molecules bound in the cavity formed π-π interaction with Y140 and hydrogen bonds with residues or water molecules around the pocket (Supplementary Fig.S7). The Hit 1 molecule at the intersection of EC2 domains bound mainly due to van der Waals interaction (Supplementary Fig. S7). The cavity around Y140 has a different structure through X dimerization from monomer and the binding mode of Hit 1 likely differs to fit the cavity.

A drastic structural change was observed in Hit 1-bound X dimer when compared with the apo form of X dimer (PDB ID; 4zmq). The angle of EC1-EC2 domain became flatter and chain B was shifted relative to chain A (Fig. 3c). This structural change in angle can be explained most reasonably by two Hit 1 molecules bound in the cavity around Y140. Hydrogen bonds including one between the side chain of K14 and the main chain of A138’ at the interface of X dimer were disrupted upon Hit 1 binding (Figure 3d), which could explain the shifted location of two monomers. The Hit 1-bound X dimer seems to be in a metastable state; therefore the Hit 1 interaction may affect the dissociation of the X dimer.

To confirm that the hydrogen bond between the side chain of K14 and the main chain of A138’ is important for X dimerization as previously reported^21^, we used a size exclusion chromatography-multiangle light scattering (SEC-MALS) analysis. Indeed, the K14A mutant was mainly in monomer form (Fig.3e). Together, these data suggest that Hit 1 binding interferes with formation of hydrogen bonds necessary for X dimerization. Unlike a typical orthosteric inhibitor, Hit 1 did not directly block hydrogen bonding between two monomer units; rather, Hit 1 alters the monomer structure to prevent hydrogen bond formation.

### Inhibition of cell adhesion by Hit 1

We next investigated whether or not Hit 1 could inhibit cell adhesion, using a previously established cell aggregation assay^33,34^. We established a CHO cell line expressing P-cadherin using the Flp-In-CHO system. In this assay, extracellular proteins other than P-cadherin are trypsinized so that cell adhesion and formation of cell aggregates depend only on the interaction of P-cadherin molecules. After the trypsinization, the culture medium was replaced with medium without calcium so that the cells were not aggregated. The aggregation reaction was initiated by addition of 1 mM CaCl_2_. As control experiments, 1 mM EDTA was added (Fig. 4a). EDTA inhibited formation of cell aggregates; thus adhesion is based on the calcium-dependent interaction of P-cadherin molecules. When Hit 1 and 1 mM CaCl_2_ were added simultaneously, aggregation was inhibited in a manner that depended on the Hit 1 concentration (Fig. 4b). When Hit 1 was added to pre-formed cell aggregates, the aggregates remained stable in the time scale of this assay; EDTA did disrupt pre-formed aggregates (Supplementary Fig. S8). These results suggest that Hit 1 does not shift the equilibrium to the monomer state at the cellular level. We hypothesize that Hit 1 blocks association of two monomers by affecting the on-rate of X dimerization.

**Fig. 4.**
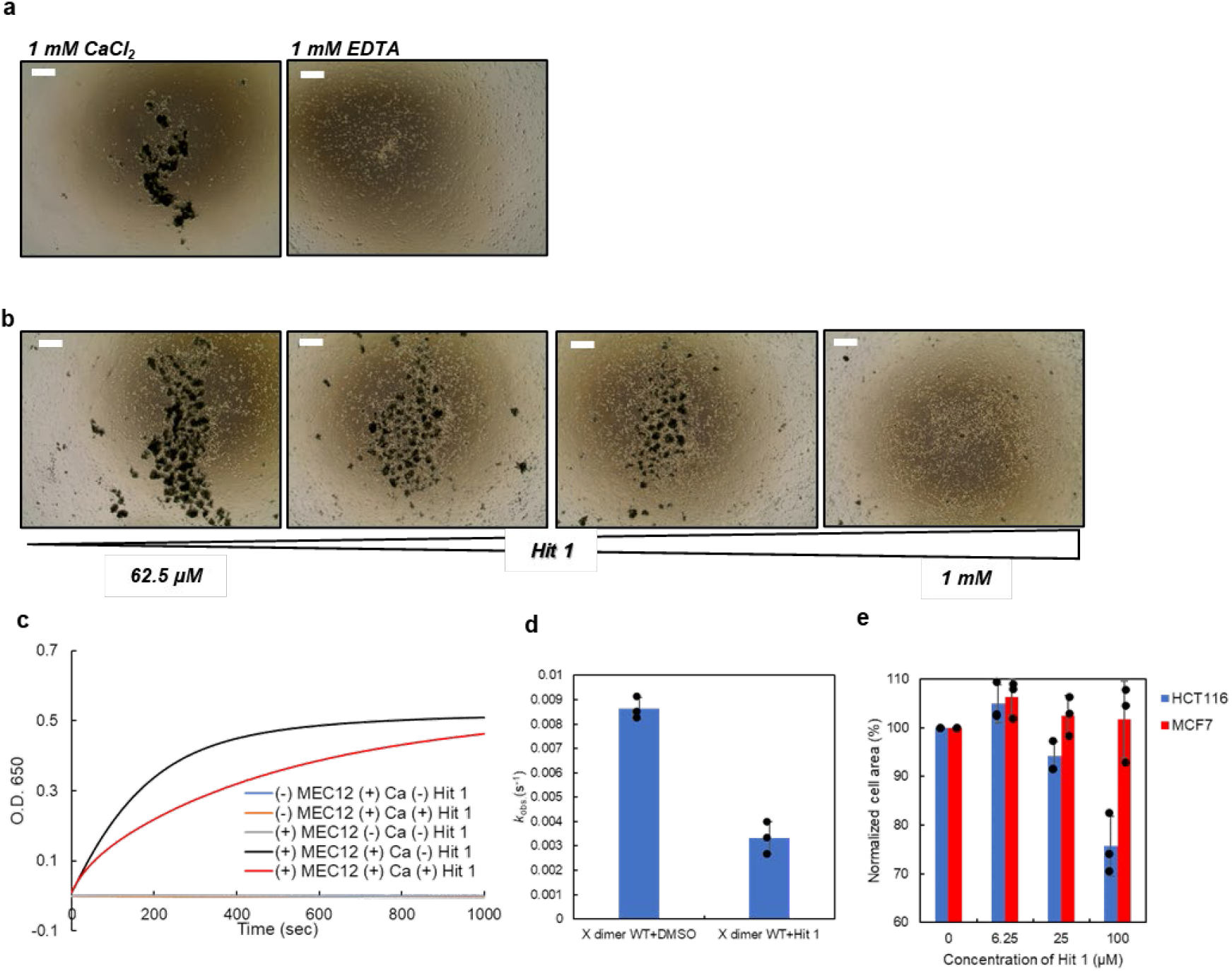
Hit 1 blocks aggregation of liposomes and cells with P-cadherin on their surfaces. (a) Images of CHO cell line expressing P-cadherin in the presence of 1 mM CaCl_2_ (left) and 1 mM EDTA (right). Scale bars indicate 500 μm. (b) Representative images of CHO cell line expressing P-cadherin in the presence of 1 mM CaCl_2_ and increasing concentrations of Hit 1. This experiment was repeated at least three times, and similar results were obtained. Scale bars indicate 500 μm. (c) Liposomes with MEC12 on the surface were incubated with CaCl_2_ and with (red) or without (black) Hit 1. Controls had no effect on optical density (blue, yellow, and grey traces). Optical density at 650 nm was monitored as a function of time. (d) The *k*_obs_, of liposome aggregation calculated assuming a one-phase association model. N=3. Error bars show standard deviation. Individual data points are shown in black plots. (e) Normalized area of HCT116 and MCF7 cells in the presence or absence Hit 1. In order to take cell shape into account, the cell area in the absence of Hit 1 was taken as 100%. N=3. Error bars show standard deviation. Individual data points are shown in black plots.

To quantify the effect of Hit 1 on the on-rate of X dimerization, we performed a liposome aggregation assay. We first prepared C terminally His-tagged MEC12 and incubated the protein with a DOPC-based liposome that contained 10% Ni-chelating lipid DOGs-NTA-Ni in the presence of EDTA to deactivate P-cadherin molecules. Under these conditions, no aggregation was observed as monitored by absorbance at 650 nm (Fig. 4c). We also confirmed that liposome aggregation only happens in the presence of both MEC12 and CaCl_2_ (Fig. 4c). Upon addition of CaCl_2_ to the solution, liposomes aggregated, presumably through X dimerization of C terminally His-tagged MEC12 molecules captured on the surface of the liposomes as indicated by a gradual increase in absorbance at 650 nm that reached a plateau (Fig. 4c). The observed rate constant (*k_obs_*) of the liposome aggregation was calculated as an indicator of the on-rate of X dimerization using an exponential decay equation model of the one-phase association. The *kobs* value was considerably lower in the presence of Hit 1 than in the absence of Hit 1 (Fig. 4d, Supplementary Fig. S9). This result supports our hypothesis that Hit 1 inhibits X dimerization and presumably cell aggregation, through on-rate modulation caused by the indirect disruption of hydrogen bonds necessary for X dimerization.

We also analyzed the effects of Hit 1 on cell adhesion mediated by endogenous P-cadherin and on cells that do not express P-cadherin using two cancer cell lines; HCT116 cells express P-cadherin^17^, and MCF7 cells do not^35^. Cells were plated in the presence of 100 μM Hit 1 and cell adhesion was quantified by cell area. The normalized cell area of HCT116 cells was significantly decreased in the presence of Hit 1 compared to its absence, but Hit 1 had no effect on normalized MCF7 cell area (Fig. 4e). This result indicates that Hit 1 blocks adhesion of cancer cells that express P-cadherin.

### Design of a more potential inhibitor

Since Hit 1 and many of the compounds identified in the primary screen had cationic functional groups, we firstly checked whether the cationic functional group is important for binding activity by testing commercially available indole-based compounds: tryptamine, tryptophan, and auxin (Fig. 5a). Surprisingly, only tryptamine bound to P-cadherin (Supplementary Fig. S10a), which may be because the protein surface around the cavity where Hit 1 binds is negatively charged (Supplementary Fig. S10b). This result indicated that the cationic functional group should not be modified during further development.

**Fig. 5.**
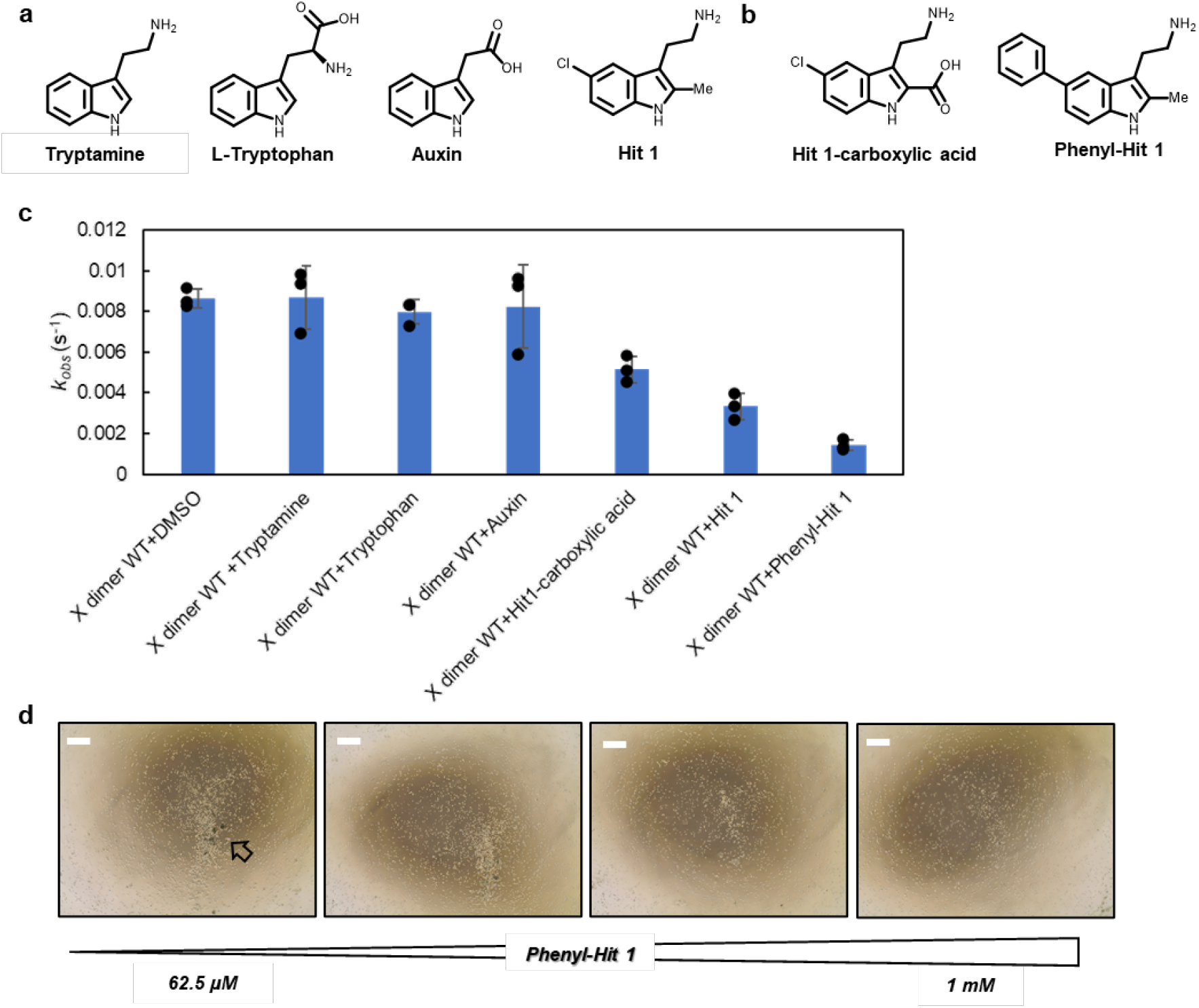
Phenyl-Hit 1 more effectively inhibits aggregation than does Hit 1. (a), (b) Chemical structures of the compounds investigated. (c) The *k*_obs_ values for indicated compounds in the liposome aggregation assay N=3. Error bars show standard deviations. Individual data points are shown in black plots. (d) Representative images from cell aggregation assay in the presence of a range of concentrations of phenyl-Hit 1. Scale bars indicate 500 μm. Black arrow indicated small aggregates formed in the presence of the lowest concentration. This experiment is repeated at least twice, and similar results were obtained.

Upon study of the 2D interaction map of Hit 1 with the P-cadherin monomer (Fig. 2c), we came up with two strategies for the further synthesis: 1) replacement of the methyl group at the second position of the indole ring with a carboxy group, which should result in formation of salt bridges between the carboxy group with R68 and the amino group with D137 and 2) replacement of the chlorine group at the C5 position of the indole ring with a bulkier functional group to inhibit approach of the second monomer. We first synthesized 3-(2-aminoethyl)-5-phenyl-1*H*-indole-2-carboxylic acid (Hit 1-carboxylic acid, Fig. 5b), but its affinity for REC12 in the SPR-based direct binding assay was no better than that of Hit 1 and it did not have significant activity in cell aggregation assay (Supplementary Fig. S11).

Next, we synthesized 2-(2-methyl-5-phenyl-1*H*-indole-3-yl)ethan-1-amine (phenyl-Hit 1) (Fig. 5b). In the SPR-based direct binding assay, we observed a dose-dependent response; the binding affinity for REC12 was equivalent to that of Hit 1 (Supplementary Fig. S12). In the liposome aggregation assay, phenyl-Hit 1 had stronger inhibitory activity than Hit 1; tryptamine auxin, and tryptophan had little effect on aggregation (Fig. 5c, Supplementary Fig. S9). Tryptamine does not have a bulky functional group, which is likely why it did not inhibit liposome aggregation. In the cell aggregation assay, phenyl-Hit 1 inhibited cell aggregation much more strongly than did Hit 1 (Fig. 5d), whereas negative controls and tryptamine did not (Supplementary Fig. S13). Given this strong inhibitory activity, the binding affinity of Phenyl-Hit 1 to X dimer could be stronger than to monomeric P-cadherin.

## Discussion

Through biophysical and structural methods, we identified a class of fragment compounds that bind to a unique shallow cavity on P-cadherin between the EC1 and EC2 domains. Our chemical fragment has a potential to modulate X dimer through the effect of on-rate modulation, by altering the angle of the EC2 domain relative to the EC1 domain, blocking formation of key hydrogen bonds necessary for X dimerization (Fig. 3c, Fig. 6). This is the first report of an inhibitor of a classical cadherin family protein that has a mechanism other than an orthosteric one. Now our fragments had an impact on kinetics; probably because the energy barrier to remove the bound ligand should be cleared to complete the X dimerization. SPR-based compound screen can provide an ideal platform to select such a kinetic modulator, since a flow system of SPR itself could help the inhibition of interaction, and thus the inhibitory mechanism should not be so strong as to be a thermodynamic one. We propose that on-rate modulation utilizing the restricted protein domain angle provides a common strategy to regulate PPI of cell adhesive molecule with a small molecule, since not only cadherin superfamily proteins but also integrin superfamily proteins and immunoglobulin superfamily proteins have multi-domain structures and similar cavities located between the domains could be targeted to regulate the protein dynamics necessary for function.

**Fig. 6.**
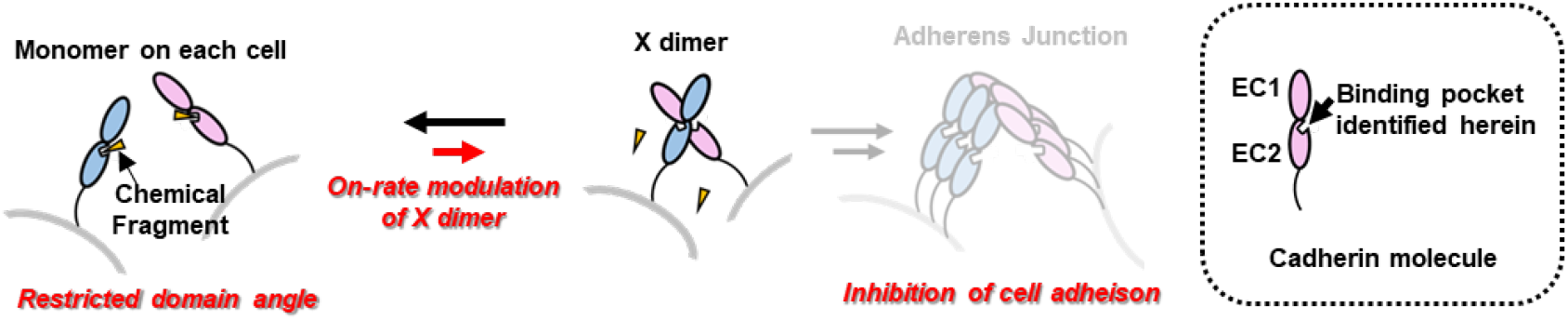
On-rate modulation mechanism of inhibition of cell adhesion by a chemical fragment. In the presence of chemical fragments, the angle between the EC1 and EC2 domains is not optimal for X dimerization; thus, the association of two monomers to form X dimer becomes slower. This kinetic effect can be strong enough to inhibit cell adhesion.

A previous study of an anti-P-cadherin single chain Fv (scFv), which thermodynamically inhibits X dimerization, indicated that thermodynamic inhibition of X dimer leads to inhibition of S-S dimerization, and thus cell adhesion^36^. Hit 1 modulated the on-rate of X dimerization, and despite the lack of a strong thermodynamic effect, inhibited cell adhesion. This result implies that every step-wise process like formation of cell adhesion assembly has a check point, such as the intermediate like X dimer state, that decreases the energy barrier to the final state and that a slight, kinetic-level inhibition of the key step can lead to significant inhibition of formation of the final state. The most effective small-molecule inhibitor of the formation of a macromolecule complex should block the initial association of the component proteins. In other words, it is this initial association of the component molecules that should be targeted for the drug discovery of the field.

From our simple structure-activity relationship study, we determined that an inhibitor with specificity for P-cadherin requires 1) a cationic functional group to interact with the negatively charged region around the cavity located between the EC1 and EC2 domains, 2) functional groups that interact with Y140, a unique residue to P-cadherin, and 3) a bulky functional group that protrudes from the interface to block X dimerization. Our structure-activity relationship also suggests that strong affinity for the protein-protein interface itself is not strictly necessary for inhibition of protein complex formation since it is not the equilibrium but rather the first contact of component molecules that an inhibitor should affect.

We discovered that tryptamine but not tryptophan binds to P-cadherin. It is known that several tryptamine-derived metabolites have their own biological role^37,38^. One tryptamine derivatives, serotonin, is important in the function of digestive organs including gastric tissue^39^; P-cadherin is overexpressed in gastric cancer. It is possible that tryptamine derivatives, through weak interactions with P-cadherin, subtly regulate cell adhesion. In support of this hypothesis, relationships between serotonin activity and cell adhesion phenotypes have been demonstrated^40–42^.

In summary, the chemical fragment we identified acts through an on-rate modulation mechanism to inhibit formation of an intermediate in dimerization of P-cadherin. Cadherin family proteins, including P-cadherin, are associated with diseases such as cancer; thus, the small-molecule regulator of dimerization we identified has therapeutic potential. Further, our strategy for kinetic regulation of protein-protein interactions and stepwise assembly of protein complexes using small molecules could be applied to identify inhibitors of formation of other macromolecular complexes.

## Methods

### Protein expression and purification

For the *in vitro* assays including SPR-based screen, crystallography, HDX-MS, and SPR-based selectivity analysis, human P-cadherin constructs and human E-cadherin constructs were expressed in *E. coli* Rosetta2(DE3). *E. coli* cells were transformed with the pET SUMO vector and used to inoculate 6 mL LB medium containing 50 mg/mL kanamycin and 34 mg/mL chloramphenicol. Cells were pre-cultured at 37 °C for 16 h then transferred into 1 L of fresh LB medium containing the same antibiotics and cultured again at 37 °C for 4 h. At this time, the O.D._600_ was around 0.5. Isopropyl *β*-D-1-thiogalactopyranoside (IPTG) was added to 0.5 mM to induce recombinant protein expression. After 16 h at 20 °C, *E. coli* cells were collected by centrifugation at 7000 g at 4 °C for 10 min, suspended in binding buffer (20 mM Tris, 300 mM NaCl, 3 mM CaCl_2_, 20 mM imidazole, pH 8.0), and sonicated for 10 min. The lysate was centrifuged at 40000 g, 4 °C, for 30 min. The supernatant was purified on a Ni-NTA agarose (QIAGEN) column, pre-equilibrated with the binding buffer. The His-tagged protein was eluted with elution buffer (20 mM Tris, 300 mM NaCl, 3 mM CaCl_2_, 300 mM imidazole, pH 8.0), and incubated in SEC buffer (10 mM HEPES, 150 mM NaCl, 3 mM CaCl_2_, pH 7.5) with Ulp1 protease to remove the SUMO protein. The resultant protein was again loaded onto a Ni-NTA agarose column and the flow-through was further purified with size-exclusion chromatography (SEC). The protein was loaded onto a Hiload 26/60 Superdex-200 column (Cytiva) pre-equilibrated with SEC buffer. All cadherin constructs were based on EC12 (1-241), which consists of two extracellular domains. For the control experiments in SPR-based direct binding screen, anti-P-cadherin scFv TSP7 was prepared. The expression host was *E. coli* BL21 (DE3). Binding buffer contained 20 mM Tris, 500 mM NaCl, 5 mM imidazole, pH 8.0. The elution buffer was 20 mM Tris, 500 mM NaCl, 300 mM imidazole, pH 8.0. In SEC process, a Hiload 26/60 Superdex-75 column (Cytiva) was used in 20 mM Tris, 200 mM NaCl, pH 8.0 buffer.

### Compounds

The fragment library used in the SPR-based screen was purchased from Drug Discovery Initiative. Hit 1 was purchased from Vitas-M Laboratory. For HDX-MS experiments, SPR-based selectivity analysis, and the mutation study, Hit 1 was synthesized in house. The synthetic scheme, procedures, and compound characterization are described in supplementary materials. Tryptamine, auxin, and L-tryptophan were purchased from Sigma Aldrich or Nacalai Tesque. Stock solution of all the compounds were prepared in DMSO and stored at −30 °C.

### Direct binding analysis using SPR

For the primary screen, we performed a direct binding assay using SPR. All the SPR-related experiments were performed on a Biacore 8K (Cytiva) at 25 °C. Monomer construct of REC12 was immobilized on the Sensor Chip SA via biotin-streptavidin capture. The immobilization level was approximately 3000 RU. Fragment compounds were injected onto the sensor chip surface in running buffer (10 mM HEPES, 150 mM NaCl, 3 mM CaCl_2_, 0.05% Tween20, 5% DMSO, pH 7.5) to a final concentration of 100 μM. Both association time and dissociation time were 20 s, and 1 M arginine-HCl, pH 4.4 was used to regenerate the sensor chip surface. The running buffer alone was used as a negative control, and 1 μM TSP7 was used as a positive control. Solvent correction was performed periodically. The binding response of each compound was normalized to those of control samples and molecular weight using the Biacore 8K software.

### ABA assay using SPR

For the secondary screen, we performed an ABA assay. EC12 was immobilized on the Sensor Chip CM5 by amine coupling (pH 4.5). The immobilization level was approximately 500 RU. Solution A was 2 μM EC12, and 100 μM compound identified in the primary screen was solution B. The association time for the first injection of solution A part was 180 s, that of solution B was 120 s, and that of the second injection of solution A was 120 s

### Selectivity analyses and mutation study

E-cadherin WT REC12, E-cadherin N140Y mutant, P-cadherin WT REC12, and P-cadherin Y140N mutant were immobilized on chips via biotin-streptavidin capture. All the constructs gave the binding response between 3000-4000 RU. Association times were 15 s or 30 s, and dissociation time was 20 s. Hit 1 concentration ranged from 37.6 μM to 500 μM. The *K_D_* values were calculated using the Scatchard method using the Biacore 8K software. Solvent correction was performed at the beginning and the end of the concentration series.

### Circular dichroism (CD) measurement

To confirm that point mutations of classical cadherin constructs did not disrupt secondary structure, we performed CD measurements. CD spectra of samples in 1 mm path-length quartz cells were measured at 20 °C using a JASCO J-820 spectropolarimeter. Samples were prepared at 10 μM in 10 mM HEPES, 150 mM NaCl, 3 mM CaCl_2_, pH 7.5 buffer.

### Crystallization of P-cadherin

For X-ray crystallography, conditions used previously for the C-terminal-deleted REC12 and MEC12 (1-213)^21^ were used. Purified C-terminal-deleted REC12 (12.5 mg/mL) was crystallized in 100 mM HEPES, pH 7.5, 28% v/v PEG 400, 200 mM CaCl_2_. Purified C-terminally-deleted MEC12 (12.5 mg/mL) was crystallized in 0.17 M sodium acetate trihydrate, 0.085 M Tris pH 8.5, w/v 25.5% PEG4000, 15% v/v glycerol. The crystals were soaked with the crystallization solution containing 10% DMSO and 10 mM Hit 1 for several minutes. For the crystals of C-terminal-deleted REC12, crystal annealing^43^ was performed in order to decrease mosaicity.

### Data collection and refinement

X-ray diffraction data sets were collected on the RIKEN Structural Genomics Beamline II (BL26B2) at SPring-8^44^. The diffraction data were processed with the KAMO programs^45–47^, and the structure was solved by molecular replacement, using the structure of P-cadherin REC12 (PDB ID 4zmz) or MEC12 (PDB ID 4zmq) as a search model with phenix.phaser. The resultant structures were iteratively refined using phenix.refine^48^ and manually rebuilt in Coot^48^. Final refinement statistics are summarized in Table 1. Figures were prepared with UCSF Chimera^49^.

### HDX-MS homodimer inhibition assay

To confirm the hypothesis from the crystallography that Hit 1 inhibits X dimerization, we performed HDX-MS and monitored the protein surface exposed to the solvent with or without Hit 1. We compared the extent to which the X dimer interface is exposed to the solvent by preparing REC12 and MEC12 at 1.5 mg/mL protein in the H2O-based 10 mM HEPES, 150 mM NaCl, 3 mM CaCl_2_, pH 7.5, 5% DMSO with or without 2 mM Hit 1. The hydrogen-deuterium exchange reaction was started by diluting D2O-based buffer by 10 fold and was quenched by addition of an equal volume of prechilled quenching buffer (8 M urea, 1 M Tris(2-carboxyethyl)phosphine hydrochloride, pH 3.0) with the HDx-3 PAL (LEAP Technologies). The quenched protein samples were subjected to online pepsin digestion and analyzed by LC-MS using UltiMate3000RSLCnano (Thermo Fisher Scientific) connected to the Q Exactive HF-X mass spectrometer (Thermo Fisher Scientific). Online pepsin digestion was performed using a Poroszyme Immobilized Pepsin Cartridge (2.1 ×30 mm; Thermo Fisher Scientific) in formic acid solution, pH 2.5 at 8 °C for 3 min at a flow rate of 50 μL/min. The desalting column and the analytical column were Acclaim PepMap300 C18 (1.0 × 15 mm; Thermo Fisher Scientific) and Hypersil Gold (1.0 × 50 mm; Thermo Fisher Scientific), respectively. The mobile phases were 0.1% formic acid solution (A buffer) and 0.1% formic acid containing 90% acetonitrile (B buffer). The deuterated peptides were eluted at a flow rate of 45 μL/min with a gradient of 10–90% of B buffer in 9 min. Mass spectrometer conditions were as follows: an electrospray voltage of 3.8 kV, positive ion mode, sheath and auxiliary nitrogen flow rate at 20 and 2 arbitrary units, ion transfer tube temperature at 275 °C, auxiliary gas heater temperature at 100 °C, and a mass range of m/z 200–2,000. Data-dependent acquisition was performed with normalized collision energy of 27 arbitrary units. The MS and MS/MS spectra were subjected to a database search analysis using the Proteome Discoverer 2.2 (Thermo Fisher Scientific) against an inhouse database containing the amino acid sequence of the C-terminal-depleted REC12. The search results and MS raw files were used for the analysis of the deuteration levels of the peptide fragments using the HDExaminer software (Sierra Analytics).

### SEC-MALS analyses

In order to measure the molecular size of each mutant, SEC-MALS analysis was performed. Purified mutants were concentrated to 3.2 mg/mL in SEC buffer and loaded onto a 10/300 Superdex 200 column (Cytiva). Size measurement was performed using a Heleos 8+ instrument (Wyatt Technology) equipped with a triple MALS/refraction index (RI)/ultraviolet detector.

### Preparation of CHO cells expressing P-cadherin

To evaluate cell adhesion, we used the Flp-In-CHO system (Life Technologies) to engineer CHO cells stably expressing full-length P-cadherin. A single clone was obtained through the limiting dilution-culture method. The expression of P-cadherin was monitored with an imaging cytometer (In Cell Analyzer 2000, Cytiva), because the DNA sequence of monomeric GFP was fused at the C-terminal of the human P-cadherin constructs. CHO cells expressing P-cadherin were cultured in Ham’s 12 medium (Life Technologies) containing 10% fetal bovine serum (FBS), 1% penicillin-streptomycin, and 0.5 mg/mL hygromycin at 37 °C, 5% CO_2_.

### Cell aggregation assay

The cell aggregation assay was performed as reported previously^33,34^. We first treated the CHO cells with 0.1% trypsin in HEPES-based magnesium-free buffer (HMF buffer, 10 mM HEPES, 137 mM NaCl, 5.4 mM KCl, 0.34 mM Na2HPO4, 1 mM CaCl_2_, 5.5 mM glucose, pH 7.4). These conditions result in digestion of protein molecules on the cell surface with the exception of P-cadherin, which is resistant to trypsinization in the presence of Ca^2+^. The trypsinization was stopped by adding HMF buffer containing 10% FBS, and trypsin was removed by washes with HMF buffer. After washing the cells with Ca^2+^-depleted HMF buffer (HCMF buffer), cells disaggregated. 500 μL of cell solution containing 1×10^5^ cells/mL was placed in wells of a 24 well plate that had been pre-treated with 1% (w/v) BSA. Addition of 1 mM CaCl_2_ in 2% DMSO initiated P-cadherin-mediated aggregation reaction. Simultaneously with CaCl_2_ addition, 1 mM EDTA in 2% DMSO as a control or Hit 1 was added. Cells were incubated at 37 °C, 80 rpm for 30 min, and images of the cells were taken with an EVOS XL Core Imaging System (Life Technologies). To determine Hit 1 could disrupt the cell aggregation, Hit 1 or EDTA was added 60 min after the addition of CaCl_2_. The images of the cells were taken after 60 min.

### Liposome aggregation assay

Lipids (DOPC: DOGs-NTA-Ni,9:1 molar ratio) were dissolved in chloroform than dried. The lipids monolayer obtained was hydrated with 10 mM HEPES, 150 mM NaCl, 3 mM CaCl_2_, pH 7.5 and subjected to 10 cycles of freeze and thaw. Liposomes were prepared using a polycarbonate filter with 100 nm pore diameter in a Mini-Extruder apparatus (Avanti). Based on a reported methodology^50^, liposomes with C-terminally His-tagged MEC12 were prepared at a lipid to protein molar ratio of 50:1 in 10 mM HEPES, 150 mM NaCl, 3 mM CaCl_2_, pH 7.5. After 10 min at room temperature an excess amount of EDTA (8.3 mM) was added to deactivate the cadherin molecules. After addition of 10 mM CaCl_2_, the optical density at 650 nm was measured using a spectrophotometer every 1 s for 1000 s. Hit 1 or phenyl-Hit 1 diluted in DMSO was added to the solution to a final DMSO concentration of 5% DMSO. The constant rate *K* was calculated using GraphPad Prism 8 software assuming exponential decay and one-phase association.

### Cell area quantification

HCT116 and MCF7 cells were detached from the plate by trypsin and resuspended to the concentration of 1×10^5^ cells/mL in McCoy’s 5 medium containing 10% FBS, and 1% penicillin-streptomycin, 0.1 % DMSO, and each concentration of Hit 1. Aliquots of 100 μL of the suspended cells were plated into wells of a 96-well plate (Greiner) and incubated at 37 °C, 5% CO_2_ for 2 days. After removing the medium, 100 μL of 10 μM Calcein AM (Invitrogen) in McCoy’s 5 medium was added to each well and incubated at 37 °C, 5% CO_2_ for 30 min. After washing with PBS buffer, 100 μL of PBS buffer was added, and images were taken using an In Cell Analyzer 2000 (Cytiva). The cell area was calculated using a method created with Developer Tool Box software. The kernel size and intensity were adjusted for each cell to fully detect the area.

## Acknowledgements

We thank Thermo Fisher Scientific for the technical support in HDX-MS experiments. This work was supported by JSPS Grants-in-Aid for Scientific Research (16H02420 to K.T.), Platform Project for Supporting Drug Discovery and Life Science Research (Basis for Supporting Innovative Drug Discovery and Life Science Research (BINDS)) from Japan Agency for Medical Research and Development (AMED) (JP19am0101094 to K.T.), and a Grant-in-Aid for JSPS fellows (A.S.). The synchrotron radiation experiments were performed at BL26B2 of SPring-8 and were supported by BINDS from AMED (JP19am0101070). The fragment compounds were provided by Drug Discovery Initiative supported by BINDS from AMED (JP19am0101086). This work was partly supported by the World-leading Innovative Graduate Study Program for Life Science and Technology, The University of Tokyo, as part of the WISE Program (Doctoral Program for World-leading Innovative & Smart Education), MEXT, Japan.

## Data Availability statements

The authors declare that the data supporting the findings of this study are available within the paper and its supplementary information files.

## Author contributions

A.S., S.N., S.K., T.T., and K.T. designed experiments, analyzed and discussed the results, and approved the manuscript. A.S., K.Y., S.I, G.U., performed crystallization experiments, and processed and determined the crystal structure. A.S. and Y.S. synthesized Hit 1, Hit 1-carboxylic acid and phenyl-Hit 1 with input from S.S.. A.S. and S.N. wrote the manuscript with input from K.T.

## Competing interests

The authors declare no competing financial interests.

## Materials & Correspondence

Supplementary information containing additional figures, and tables and compound characterization data is available as a separate file. Correspondence should be addressed to K.T. or S.N.

## Notes

### Competing Interest Statement

The authors have declared no competing interest.

