## Supplementary information for "On-rate modulation of cadherin interactions by chemical fragments"

For

Senoo *et al.*

**Supplementary Figures**

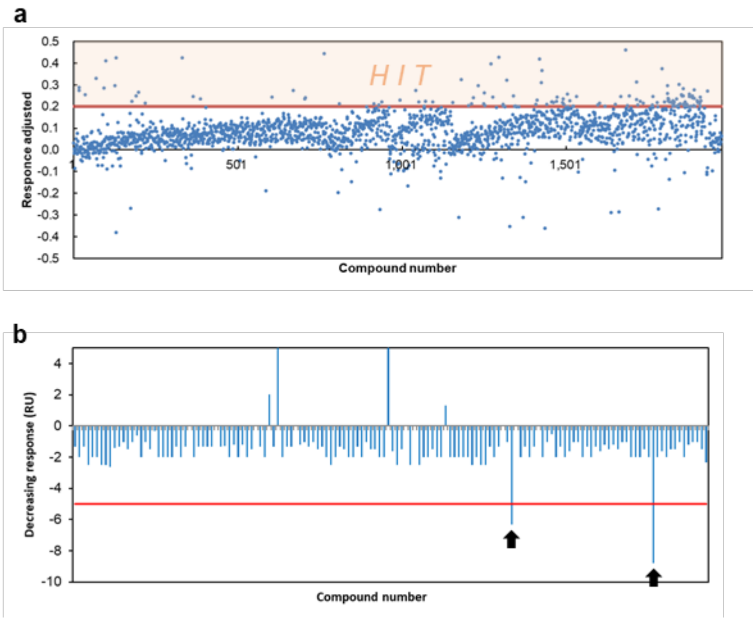


**Supplementary Fig. S1 | SPR-based fragment screen.** (a) Results of primary direct binding screen of 1973 compounds; compounds with adjusted response greater than 0.2 were regarded as hits. Adjusted response level was determined based on the response of the positive control, TSP7 (anti-P-cadherin scFv), and the compound molecular weight. The adjusted response of 0.2 is approximately half of $R_{MAX}$. (b) Results of secondary ABA screen. Compounds with a significant decreased response to addition of solution B were selected. These compounds caused dissociation of a monomer not covalently linked to the sensor chip.

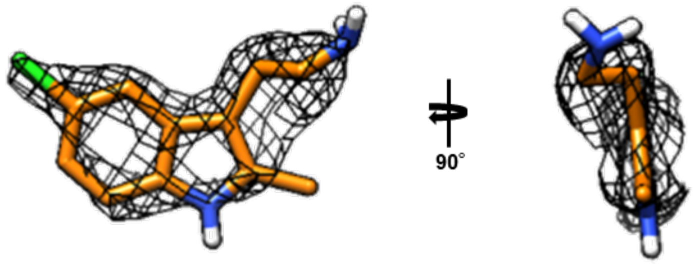


**Supplementary Fig. S2 | 2mFo-DFc map of Hit 1.** 2mFo-DFc map for Hit 1 from REC12-Hit 1 complex at 1.0 σ level is shown.

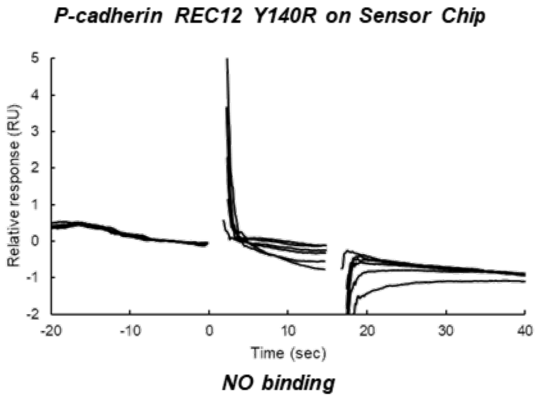


**Supplementary Fig. S3 | Hit 1 does not bind to REC12 Y140R.**

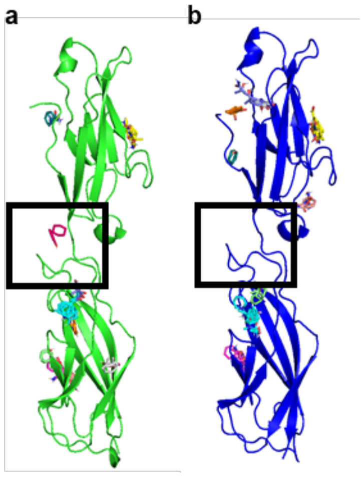


**Supplementary Fig. S4 | Binding cavity is detected in P-cadherin, but not in E-cadherin.** FTMaps of (a) P-cadherin REC12, and (b) E-cadherin EC12. Black square indicates the regions of interest.

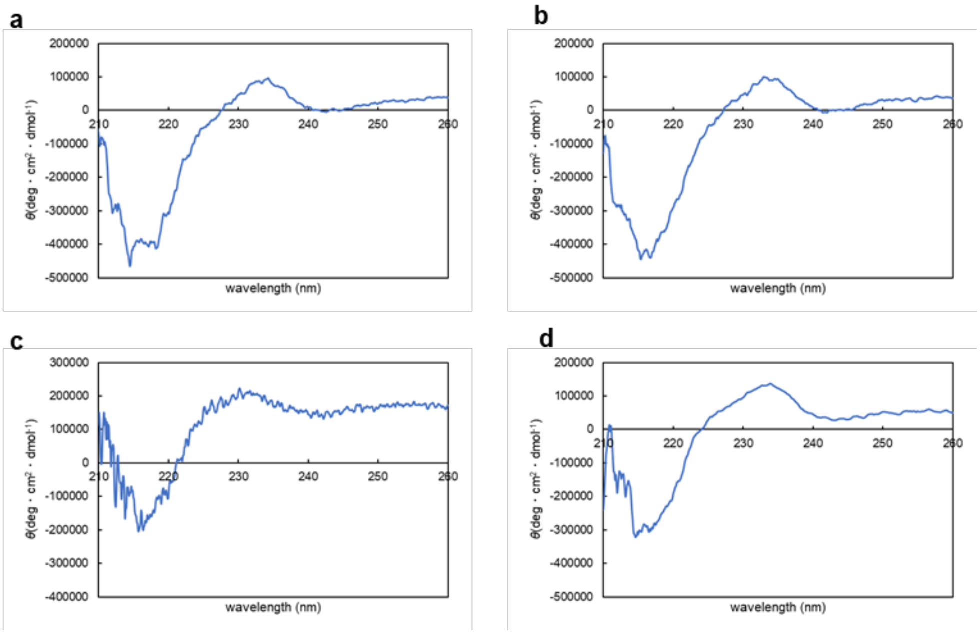


**Supplementary Fig. S5 | Secondary structure of P-cadherin REC12 WT and Y140R, and E-cadherin REC12 WT and N140Y.** Circular dichroism spectra of (a) P-cadherin REC12 WT, (b) P-cadherin REC12 Y140R, (c) E-cadherin REC12 WT, (d) E-cadherin REC12 N140Y.

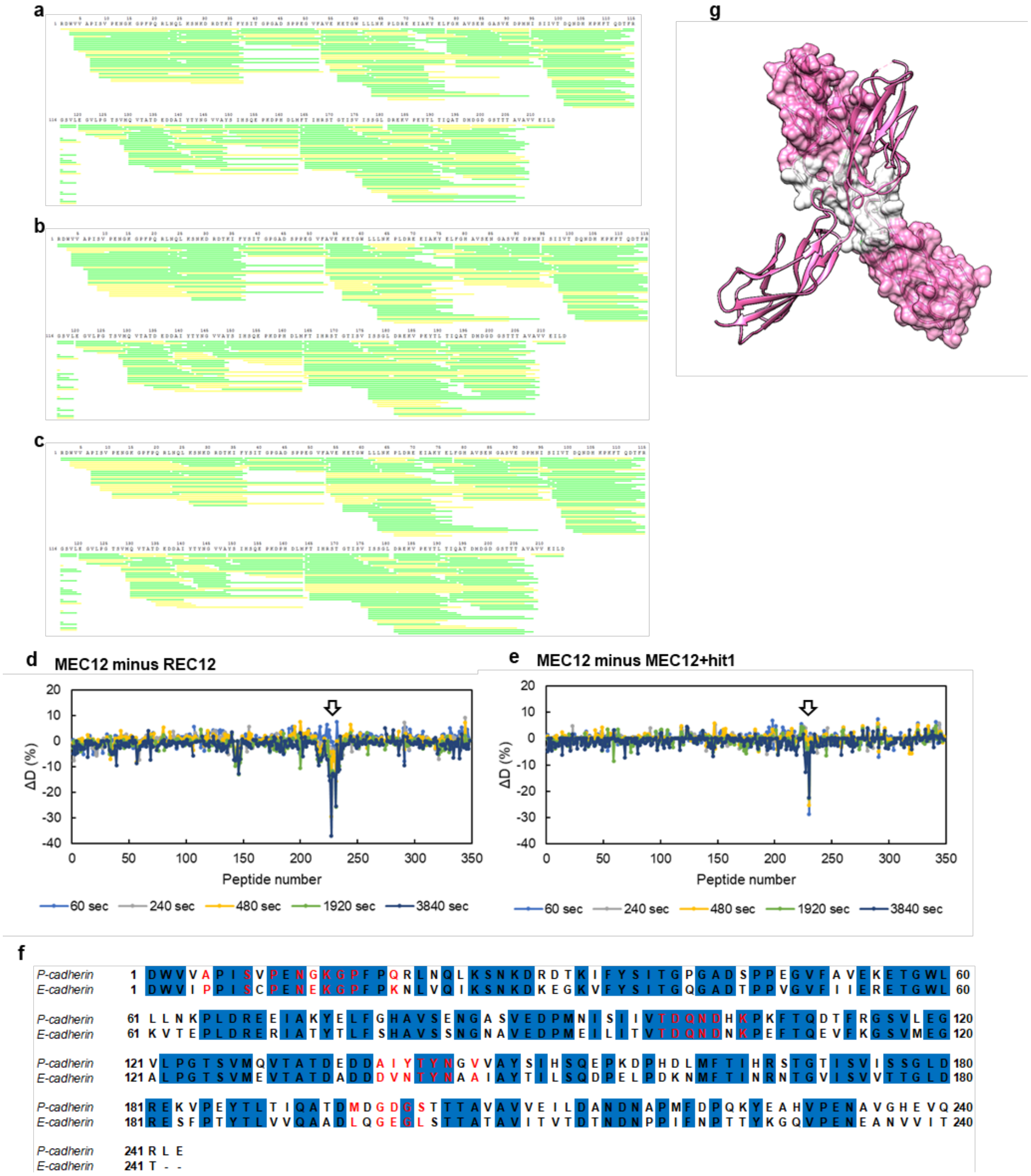


**Supplementary Fig. S6 | Hydrogen-deuterium exchange mass spectrometry as assay for X dimerization.** (a-c) Each bar represents a peptide in a) MEC12, b) MEC12 with Hit 1, and c) REC12. In all the samples, the whole region of the protein was covered in HDX-MS. Yellow and green indicate whether the confidence in the peptide identified is medium or high, respectively. (d, e) Residual plots of hydrogen-deuterium exchange ratio at each peptide. That of MEC12 minus that of REC12 is shown in d), and that of MEC12 minus that of MEC12 with Hit 1 is shown in e). The black arrow indicates the peptide region where the deuterium exchange rate was higher d) in monomer than in X dimer or e) in the X dimer with Hit 1 than the X dimer without Hit 1. (f) Sequence alignment between P-cadherin and E-cadherin. Blue background represents identical residues. Red characters represent the interface of X dimer. (g) Structure of MEC12, X dimer construct (PDB ID; 4zmq). One of the monomers is shown in surface. White region represents the interface of X dimer.

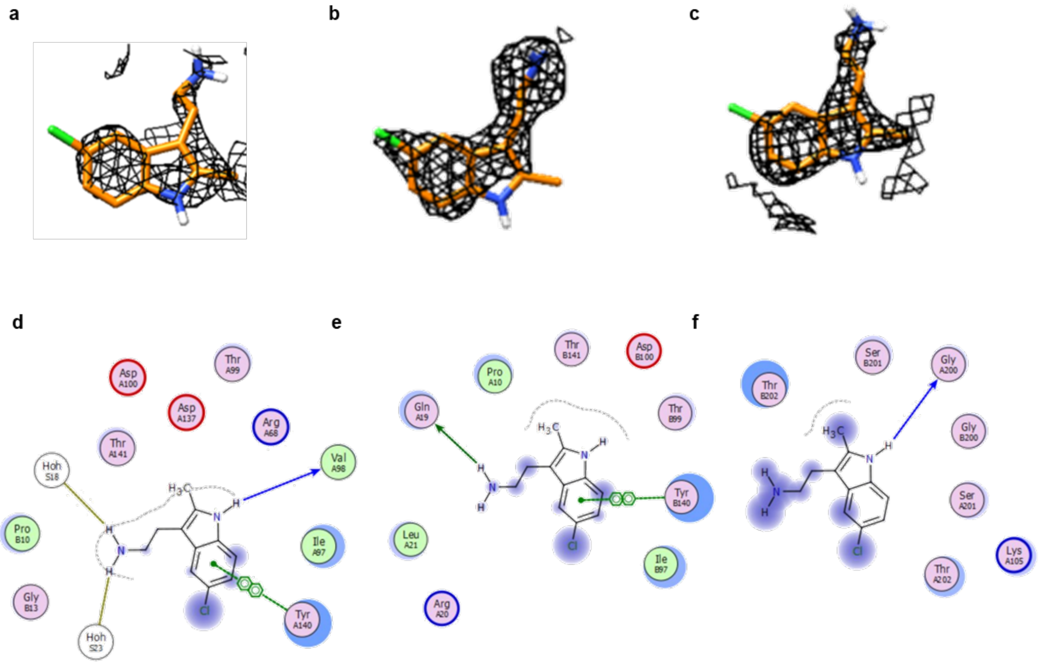


**Supplementary Fig. S7 | Structure of complex between MEC12 and Hit 1.** (a-c) 2mFo-DFc map at 0.9 $\sigma$ level for Hit 1 associated with a) chain A, b) chain B, and c) the intersection of EC2. (d-f) 2D interaction maps of Hit 1 associated with d) chain A, e) chain B, and f) the intersection of EC2 domain made with FLEV. Green arrow shows contact with side chain acceptor. Green dotted line shows $\pi$-$\pi$ interaction. Blue arrow shows contact with backbone acceptor or backbone donor. Goldenrod line shows contact with solvent water.

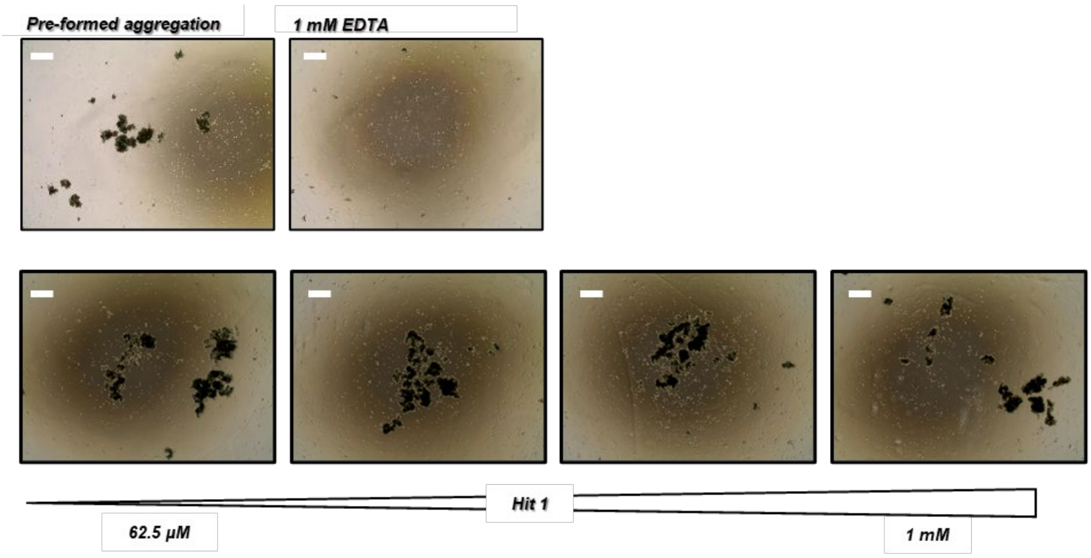


**Supplementary Fig. S8 | Hit 1 does not disrupt cell aggregates.** Upper: Images of pre-formed cell aggregates and a sample of pre-formed cell aggregates treated with 1 mM EDTA. Lower: Images of pre-formed cell aggregates treated with increasing concentrations of Hit 1. Photographs were taken 60 min after EDTA or Hit 1 addition. Scale bars indicate 500 µm.

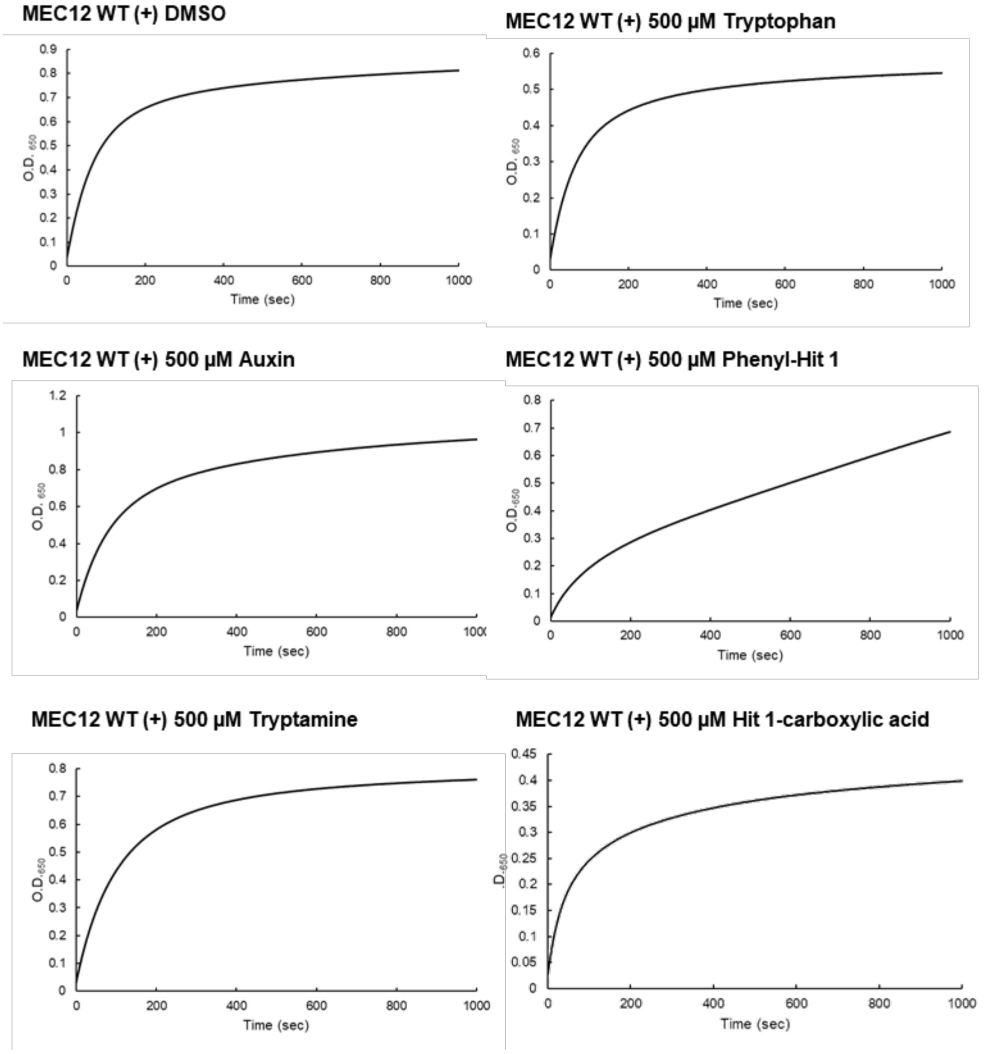


**Supplementary Fig. S9 | Liposome aggregation assays with indole derivatives, phenyl-Hit 1, and Hit 1-carboxylic acid.** Time course of optical density at 650 nm. The rate constant values in Fig. 4d and Fig. 5c were calculated using these raw data.

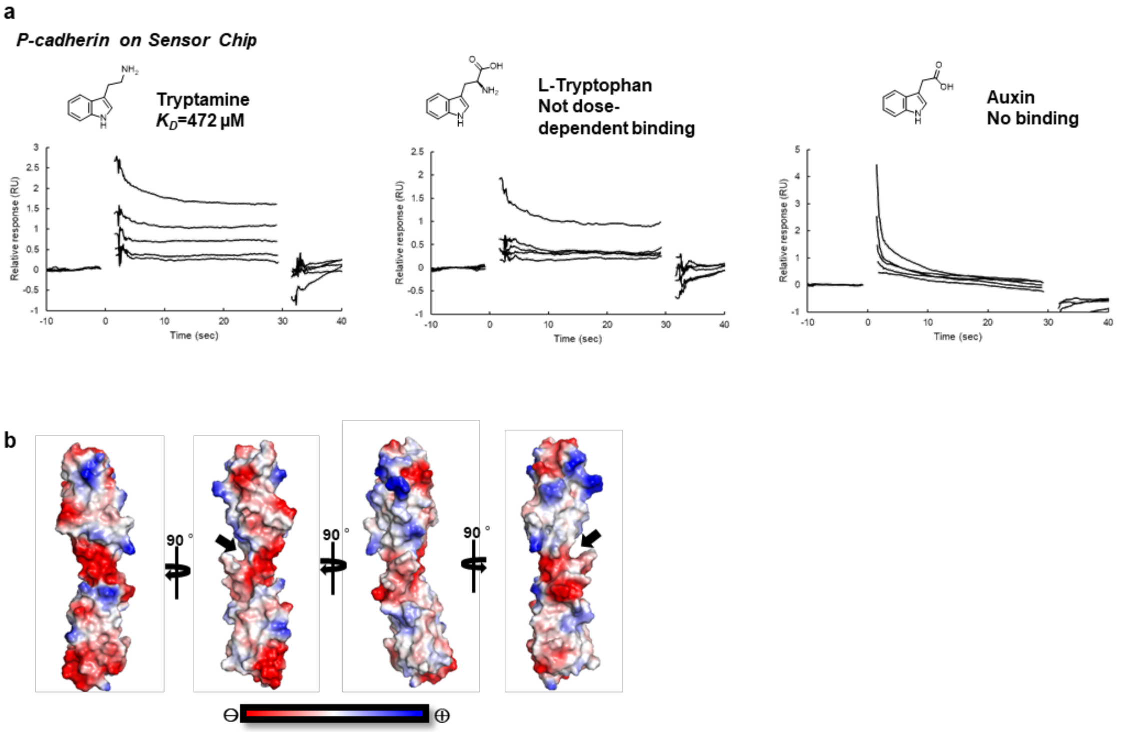


**Supplementary Fig. S10 | Tryptamine binds to P-cadherin REC12.** (a) Sensorgrams of tryptamine, auxin, and L-tryptophan injected onto immobilized P-cadherin REC12. (b) Surface representation of electrostatic potential of REC12 (PDB ID; 4zmz). The electrostatic gradient from red to blue represents -73 to +73 kT. The binding cavity is indicated by a black arrow.

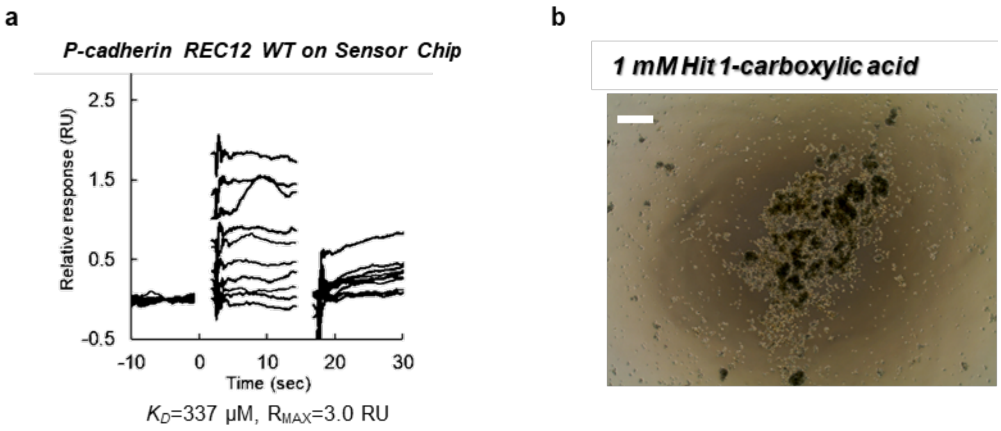


**Supplementary Fig. S11 | Characterization of activities of Hit 1-carboxylic acid.** (a) The binding of Hit 1-carboxylic acid on P-cadherin by SPR. P-cadherin REC12 was immobilized on the Sensor Chip SA and Hit 1-carboxylic acid was injected. Hit 1-carboxylic acid concentration ranged from 37.6 μM to 500 μM. (b) Image of cells incubated with 1mM Hit 1-carboxylic acid. Scale bar indicates 500 μm.

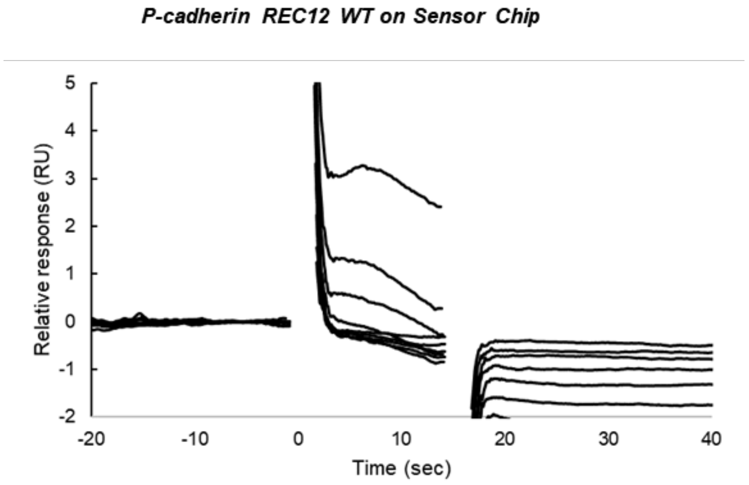


**Supplementary Fig. S12 | SPR demonstrates phenyl-Hit 1 binding to P-cadherin.** P-cadherin REC12 was immobilized on the Sensor Chip SA, and phenyl-Hit 1 was injected. Concentration of phenyl-Hit 1 ranged from 37.6 μM to 500 μM.

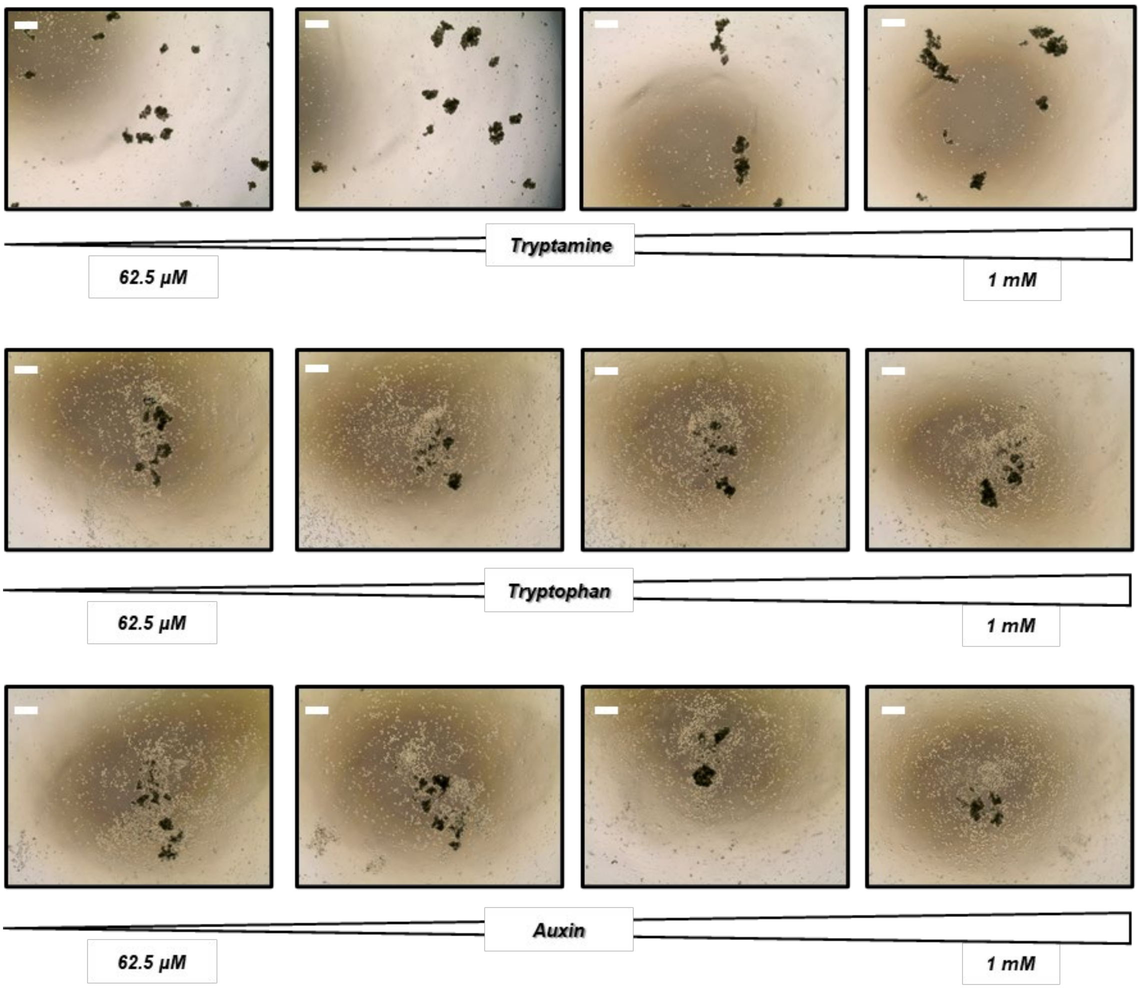


**Supplementary Fig. S13 | Tryptamine, tryptophan, and auxin do not inhibit cell aggregation.** Images of cells incubated with the indicated range of concentrations of tryptamine, tryptophan, and auxin. Scale bar indicates 500 μm.

### Supplementary methods

#### General

All reagents and dry solvents were purchased from commercial suppliers and used without further purification. Nuclear magnetic resonance (NMR) spectra were recorded using a JEOL ECS400 ($^{1}H$ 400 MHz, $^{13}C$ 100 MHz) spectrometer. Chemical shifts for $^{1}H$ NMR are expressed in parts per million (ppm) relative to tetramethylsilane (δ 0.00 ppm) in $CDCl_3$, $CHD_2OD$ (δ 3.31 ppm) in $CD_3OD$, HOD (δ 4.79 ppm) in $D_2O$, and DMSO-$d_5$ (δ 2.50 ppm) in DMSO-$d_6$. Chemical shifts for $^{13}C$ NMR are expressed in parts per million (ppm) relative to $CDCl_3$ (δ 77.0 ppm), $CD_3OD$ (δ 49.0 ppm), DMSO-$d_6$ (δ 39.5 ppm), and 1,4-dioxane (δ 67.2 ppm, in $D_2O$). Data are reported as follows: chemical shift, multiplicity (s = singlet, brs = broad singlet, d = doublet, dd = doublet of doublets, t = triplet, m = multiplet), coupling constant (Hz), and integration. High-resolution mass spectra (HRMS) were measured using a Bruker micrOTOF II mass spectrometer (Bruker ESI-TOF) and a JEOL JMS-T100LP AccuTOF LC-plus (JEOL ESI-TOF) mass spectrometer.

**Synthesis of *tert*-butyl(2-(5-bromo-2-methyl-1*H*-indole-3-yl)ethyl)carbamate**

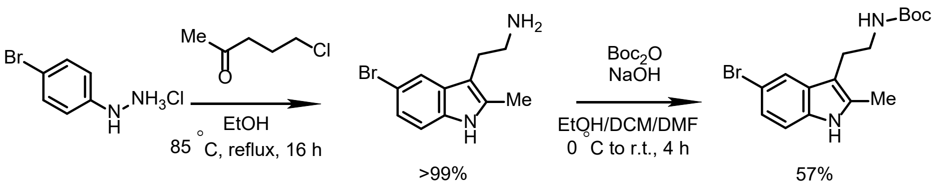


To a solution of 4-bromo-phenylhydrazine (149 mg, 667 μmol) in EtOH (2.0 mL) was added 5-chloropentan-2-one (150 μL, 1.31 mmol, 2.0 equiv.). The reaction mixture was stirred at 85 °C for 16 h. Then, the solvent was removed under reduced pressure. To the resultant solution of 2-(5-bromo-2-methyl-1*H*-indol-3-yl)ethan-1-amine was added two pellets of NaOH (*ca.* 180 mg, *ca*. 4.5 mmol), and then added dropwise a solution of di-*tert*-butyl dicarbonate (200 μL, 871 μmol, 0.9 equiv.) in EtOAc/DCM/DMF (2.0 mL/2.0 mL/2.0 mL) at 0 °C. The mixture was warmed to room temperature and stirred for 4 h. Then the reaction mixture was quenched with $NaHCO_3$ saturated solution and extracted with ethyl acetate. The combined organic layers were washed with brine, dried over $Na_2SO_4$ and filtrated. After the solvent was removed under reduced pressure, the residue was purified by silica gel column chromatography with Hexane-Ethyl acetate (96%-4% to 60%-40%) to give compound as blown foam (141 mg, 57%). $^{1}H$ NMR ($CDCl_3$, 400 MHz) $\delta$ = 1.36 (s, 9H), 2.24 (s, 3H), 2.73 (t, $J$ = 6.4 Hz, 2H), 3.22–3.24 (m, 2H), 4.55 (brs, 1H), 7.01 (d, $J$ = 8.4 Hz, 1H), 7.08 (dd, $J$ = 8.4, 1.6 Hz, 1H), 7.49 (s, 1H), 8.20 (brs, 1H); $^{13}C$ NMR ($CDCl_3$, 100 MHz) $\delta$ = 11.5, 24.5, 28.4, 41.1, 49.2, 108.3, 111.7, 112.4, 120.3, 123.6, 130.4, 133.5, 133.9, 156.0; HRMS (JEOL ESI-TOF): *m*/*z* calcd. for $C_{16}H_{21}BrN_2NaO_2^+$ [M+Na]$^+$ = 377.0664, found = 377.0647.

**Synthesis of *tert*-butyl(2-(5-chloro-2-methyl-1*H*-indole-3-yl)ethyl)carbamate**

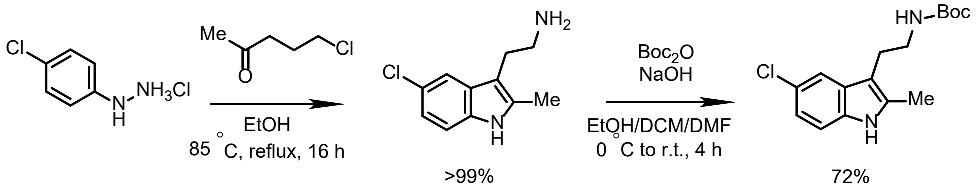


The titled compound was synthesized according to the procedure described above, using 4-chlorophenylhydrazine hydrochloride instead of 4-bromophenylhydrazine hydrochloride. The yield was 72%. $^{1}H$ NMR ($CDCl_3$, 400 MHz) $\delta$ = 1.36 (s, 9H), 2.25 (s, 3H), 2.74 (t, $J$ = 6.4 Hz, 2H), 3.22–3.24 (m, 2H), 4.52 (brs, 1H), 6.95 (d, $J$ = 8.8 Hz, 1H), 7.06 (d, $J$ = 8.8 Hz, 1H), 7.34 (s, 1H), 8.08 (brs, 1H); $^{13}C$ NMR ($CDCl_3$, 100 MHz) $\delta$ = 11.5, 24.5, 28.4, 41.0, 79.2, 108.5, 111.2, 117.3, 121.1, 124.9, 129.8, 133.6, 133.7, 156.0; HRMS (Burker-ESI): *m*/*z* calcd. for $C_{16}H_{21}ClN_2NaO_2^+$ $[M+Na]^+$ = 331.1184, found = 331.1184.

**Synthesis of 2-(5-chloro-2-methyl-1*H*-indole-3-yl)ethan-1-amine (Hit 1)**

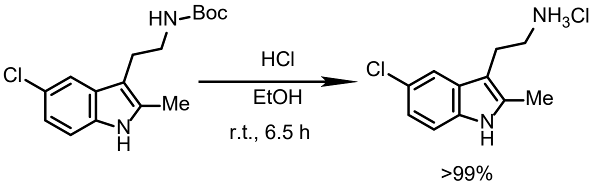


To a screw tube containing *tert*-butyl(2-(5-chloro-2-methyl-1*H*-indole-3-yl)ethyl)carbamate (100 mg, 323 μmol) was added 4 M HCl in ethyl acetate (1.0 mL). The reaction mixture was stirred at room temperature for 6.5 h. The solvent was removed under reduced pressure to give Hit 1 as a hydrogen chloride salt, brownish powder (81.1 mg, >99%). $^{1}H$ NMR ($D_2O$, 400 MHz) $\delta$ = 2.41 (s,

3H), 3.08 (t, $J$ = 6.4 Hz, 2H), 3.26 (t, $J$ = 6.8 Hz, 2H), 7.17 (d, $J$ = 8.4 Hz, 1H), 7.40 (d, $J$ = 8.4 Hz, 1H), 7.58 (s, 1H); $^{13}$C NMR ($D_2O$, 100 MHz) $\delta$ = 11.1, 22.0, 40.2, 105.0, 112.7, 117.1, 121.4, 124.8, 129.3, 134.3, 136.7; HRMS (Bruker ESI-TOF): $m/z$ calcd. for $C_{11}H_{15}Cl_2N_2^+$ $[M+H]^+$ = 209.0840, found = 209.0849.

**Synthesis of 2-(2-methyl-5-phenyl-1*H*-indole-3-yl)ethan-1-amine (phenyl-Hit 1)**

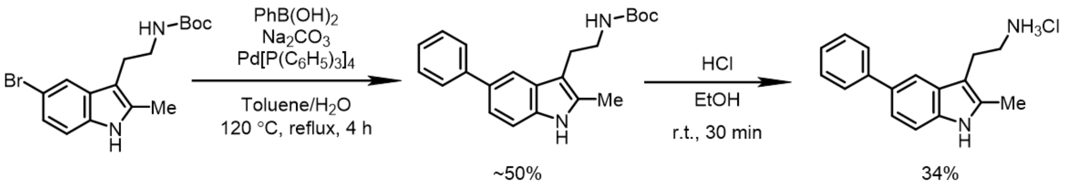


To a screw tube were added *tert*-butyl(2-(5-bromo-2-methyl-1*H*-indole-3-yl)ethyl)carbamate (76.3 mg, 216 μmol), phenylboronic acid (52.7 mg, 432 μmol, 2.0 equiv.), sodium carbonate (59.7 mg, 563 μmol, 2.6 equiv.) and $Pd[P(C_6H_5)_3]_4$ (25.0 mg, 10 mol%), and the tube was filled with nitrogen gas. To the mixture, toluene (1.1 mL) and degassed water (220 μL) were added and the mixture was stirred at reflux for 4 h. The mixture was diluted and extracted with ethyl acetate. The combined organic layers were washed with brine, dried over $Na_2SO_4$ and filtrated. The residue was purified by silica gel column chromatography with Hexane-Ethyl acetate (96%-4% to 60%-40%) to give brown foam. Because the desired product and remained substrate were inseparable, the mixture of these two compounds were used for the next step. To another screw tube were added the mixture of compounds (27.4 mg) and 4 M HCl in ethyl acetate (500 μL). The reaction mixture was stirred at room temperature for 30 min. The resultant compound was purified using HPLC with water (0.1% TFA)-Acetonitrile (0.1% TFA) (70%-30% to 30%-70%) to give Phenyl-Hit 1 as a salt of trifluoroacetic acid, white powder (9.2 mg, 34%). $^{1}$H NMR (methanol-$d_4$, 400 MHz) $\delta$ = 2.43 (s, 3H), 3.11–3.17 (m, 4H), 7.24–7.41 (m, 5H), 7.63–7.69 (m, 3H); $^{13}$C NMR (methanol-$d_4$, 100 MHz) $\delta$ = 11.3, 23.4, 41.3, 106.3, 111.9, 116.4, 121.5, 127.1, 128.1, 129.6, 129.9, 133.8, 135.0, 136.9, 144.2; HRMS (Bruker ESI-TOF): *m/z* calcd. for $C_{17}H_{19}N_2^+$ $[M+H]^+$ = 251.1543, found = 251.1547.

**Synthesis of 6-chloro-2,3,4,9-tetrahydro-1*H*-pyrido[3,4-b]indol-1-one**

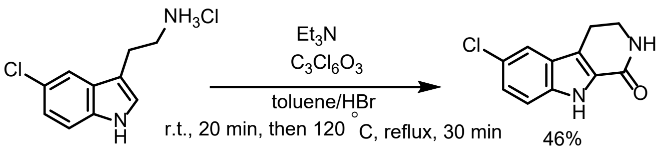


To a solution of 2-(5-chloro-1*H*-indol-3-yl)ethan-1-amine (201 mg, 870 μmol) in toluene (15 mL) were added triethylamine (140 μL, 472 μmol, 0.54 equiv.) and bis(trichloromethyl) carbonate (212 mg, 2.09 mmol, 2.4 equiv.) by stirring vigorously. The reaction mixture was stirred for 20 min at room temperature. Then 590 μL of hydrogen bromide was added to the solution, and the reaction mixture was stirred at reflux for 30 min. After cooling the solution to room temperature, water was added to quench the reaction. The reaction mixture was extracted with ethyl acetate. The combined organic layers were washed with brine, dried over $Na_2SO_4$ and filtrated. After the solvent was removed under reduced pressure, the residue was purified by silica gel column chromatography with Hexane-Ethyl acetate (60%-40% to 100%-0%) to give the desired product as a yellowish powder (87.0 mg, 46%). $^{1}$H NMR (methanol-$d_4$, 400 MHz) $\delta$ = 2.99 (t, $J$ = 7.2 Hz, 2H), 3.63 (t, $J$ = 7.2 Hz, 2H), 7.21 (d, $J$ = 8.8 Hz, 1H), 7.41 (d, $J$ = 9.2 Hz, 1H), 7.58 (s, 1H); $^{13}$C NMR (methanol-$d_4$, 100 MHz) $\delta$ = 21.4, 42.7, 114.8, 120.4, 120.5, 126.0, 126.6, 127.4, 129.0, 137.4, 136.1, 164.5; HRMS (Bruker ESI-TOF): *m*/*z* calcd. for $C_{11}H_8ClN_2O^-$ [M-H]$^-$ = 219.0331, found = 219.0338.

**Synthesis of 3-(2-aminoethyl)-5-chloro-1*H*-indole-2-carboxylic acid (Hit 1-carboxylic acid)**

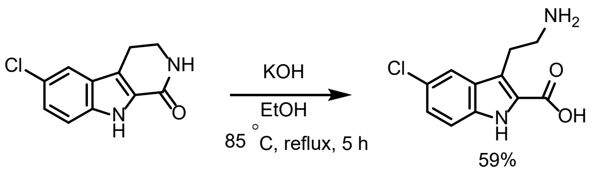


To powder of 6-chloro-2,3,4,9-tetrahydro-1*H*-pyrido[3,4-b]indol-1-one (18.8 mg, 85.1 μmol), 2 M KOH in $H_2O$/EtOH (2.0 mL/2.0 mL) was added and the reaction mixture was stirred at 85 °C for 5 h under reflux condition. The reaction mixture was cooled to room temperature and the solvent was removed under reduced pressure. To the reaction mixture was added acetic acid until the white powder dissolved. Acetic acid was removed under reduced pressure to give Hit 1-carboxylic acid as an acetate, white powder (15.0 mg, 59%). $^{1}$H NMR (DMSO-$d_6$, 400 MHz) $\delta$ = 2.97–3.01 (m, 2H), 3.16–3.19 (m, 2H), 7.07 (d, $J$ = 8.8 Hz, 1H), 7.33 (d, $J$ = 8.8 Hz, 1H), 7.61 (s, 1H), 11.1 (brs, 1H); $^{13}$C NMR (DMSO-$d_6$, 100 MHz) $\delta$ = 22.7, 40.4, 110.2, 113.3, 118.0, 121.9, 122.9, 128.9, 133.0, 136.5, 165.1; HRMS (Bruker ESI-TOF): *m*/*z* calcd. for $C_{11}H_{12}ClN_2O^+$ [M+H]$^+$ = 239.0582, found = 239.0584.

**Supplementary Table 1 | Crystallographic data collection and refinement statistics**

| Data collection | REC12-Hit 1 | MEC12-Hit 1 |
|---|---|---|
| **PDB ID** | 7CMF | 7CME |
| **space group** | *C* 1 2 1 | $P\ 2_1\ 2_1\ 2_1$ |
| **unit cell dimensions** | | |
| *a, b, c* (Å) | 74.3 40.8 72.6 | 79.8 99.1 107.9 |
| *α, β, γ* (°) | 90 97.4 90 | 90 90 90 |
| wavelength (Å) | 1.0000 | 1.0000 |
| resolution (Å)* | 36.85 - 2.30 (2.39 - 2.30) | 45.04 - 2.45 (2.60 - 2.45) |
| $R_{\mathrm{merge}}$ | 0.15 (1.09) | 0.07 (1.56) |
| $R_{meas}$ | 0.19 (1.28) | 0.08 (1.67) |
| $CC_{1/2}$ | 0.99 (0.53) | 0.99 (0.54) |
| $\langle I/\sigma(I)\rangle$ | 7.80 (1.28) | 19.21 (1.30) |
| completeness (%) | 94.4 (97.3) | 99.9 (99.8) |
| redundancy | 3.1 (3.1) | 7.4 (7.5) |
| **Refinement statistics** | | |
| resolution (Å) | 36.85-2.30 | 45.04–2.45 |
| $R_{\mathrm{work}}$ | 0.245 | 0.209 |
| $R_{\mathrm{free}}$ | 0.287 | 0.249 |
| No. of non-hydrogen atoms | 1664 | 3388 |
| macromolecules | 1626 | 3303 |
| ligands | 18 | 56 |
| solvent | 20 | 29 |
| unique reflections | 9268 (940) | 31974 (3199) |
| Average B-factor (Å$^2$) | 54.57 | 75.81 |
| **R. M. S. deviations from ideal** | | |
| bonds (Å) | 0.005 | 0.01 |
| angles (°) | 0.77 | 1.15 |
| **Ramachandran plot (%)** | | |
| Favored region | 94.76 | 96.46 |
| Allowed region | 4.76 | 3.07 |
| Outlier region | 0.48 | 0.47 |

* Values in parentheses are for the highest-resolution shell.
